# SETD6-mediated methylation of PPARγ establishes a transcriptional feedback circuit promoting lipid accumulation in liver-derived cells

**DOI:** 10.64898/2026.04.10.717761

**Authors:** Noa Nashnaz, Dana Goldberg, Maayan Abramov, Anand Chopra, Tamar Rosiecki, Habib Muallem, Yulia Haim, Tzofit Elbaz Biton, Liron Levin, Raz Zarivach, Michal Feldman, Assaf Rudich, Dan Levy

## Abstract

Peroxisome proliferator-activated receptor gamma (PPARγ) is a key transcriptional regulator of genes mediating adipogenesis (fat-cell differentiation) and lipid storage in several cell types like hepatocytes. As such, its regulation is crucial for cell and organismal physiology. Indeed, PPARγ’s activity is regulated by multiple mechanisms, including post-transcriptional modifications, which, when dys-coordinated, may contribute to the pathogenesis of various states, including obesity, insulin resistance, and fatty liver disease. Here, we demonstrate that SETD6 binds to, and methylates PPARγ at lysine 170 (K170) both *in vitro* and in liver-derived cells. This methylation event, in turn, is required for PPARγ-mediated activation of *SETD6* transcription via promoter binding, forming a positive feedback regulatory loop. RNA sequencing revealed that both SETD6 and PPARγ methylation at K170 are required for full induction of lipid metabolism genes’ expression, manifesting functionally in lipid droplet biogenesis in liver-derived cells. Together, our findings uncover a novel role for lysine methylation of PPARγ in the regulation of lipid synthesis and lipid droplet biogenesis, thereby identifying putative new therapeutic targets for lipid over-production diseases, including MAFLD (Metabolic dysfunction-associated fatty liver disease) and obesity.

## Introduction

PPAR family members are ligand-activated nuclear receptors that function as transcription factors. The family consists of three PPAR isotypes: PPARα, PPARβ/δ, and PPARγ, with highly conserved DNA and ligand binding domains (1,2). The DNA binding domain enables the binding to consensus DNA sequences called peroxisome proliferator response elements (PPREs), usually found in the promoter region of genes (3). Despite their similar domain structure and mechanism of action, PPARs are encoded by different genes and activated by different ligands. PPARα is highly expressed in oxidative tissues, such as the liver, skeletal muscle, brown adipose tissue, heart, and kidney (4). It mainly participates in the fasting state and regulates the transcription of rate-limiting enzymes required for peroxisomal and mitochondrial beta-oxidation (4). PPARβ/δ is ubiquitously expressed and has mainly been studied in skeletal muscle (5). Similar to PPARα, it has been shown to have anti-inflammatory effects (6). PPARγ has two main isoforms – PPARγ1 and PPARγ2 (7). PPARγ — and specifically PPARγ2 (from here on referred to as PPARγ) — appears to be a major driver of adipogenesis (fat cell differentiation)(8,9), and is the major isoform expressed in hepatocytes, driving transcriptional programs for fatty acid uptake, lipogenesis and lipid droplet biogenesis

Lysine methylation, among other well-studied post-translational modifications (PTMs), is a critical player in the regulation of many cellular signaling pathways. The methylation of lysine residues is performed by protein lysine (K) methyltransferases (PKMTs), which can selectively catalyze mono-, di-, or tri-methylation (10). There are over 60 members of this enzyme family, the vast majority of which contain a conserved SET domain responsible for the enzymatic activity (10,11). The mono-methyltransferase SET domain–containing protein 6 (SETD6) catalyzes the methylation of various non-histone proteins involved in regulating, among others, the NFκB pathway, oxidative stress response, WNT signaling, cell cycle, and embryonic stem cell self-renewal (12). Many of these pathways and processes have been linked to metabolism in general, and to fatty liver diseases. However, the role of lysine methylation and protein lysine methyltransferases (PKMTs) in the regulation of non-histone proteins in lipid droplet formation and steatosis remains largely unexplored.

Here, we uncover a new role of SETD6 in the accumulation of lipid droplets in human liver-derived cells. Our findings show that SETD6 binds and methylates PPARγ on lysine K170 at chromatin. PPARγK170 methylation positively regulates the formation of lipid droplets in liver cells through promoting PPARγ binding to its target gene promotors, thereby selectively activating genes involved in lipid droplet formation. Additionally, we demonstrate a functional positive feedback loop mechanism, in which PPARγ regulates the expression of SETD6 mRNA via a SETD6 and in a PPARγ K170 methylation-dependent manner. Together, these results highlight a novel pathway by which SETD6 contributes to steatosis progression through the methylation of PPARγ, positioning SETD6 as a critical regulator of lipid metabolism and presumably of the development of Metabolic dysfunction-associated fatty liver disease (MAFLD).

## Results

### *SETD6* is a target gene of PPARγ in HepG2 cells

PPARγ has previously been shown to bind to a PPRE in the *SET7/9* promotor and enhance its transcription (13). Interestingly, using the JASPAR CORE database (14), we identified a significant predicted Peroxisome Proliferator Response Element (PPRE) also within the promoter region of *SETD6* (**Fig. 1A**). This putative PPRE binding site implies that PPARγ may regulate SETD6 expression through binding to the SETD6 promoter. Analysis of publicly available ChIP-sequencing experiments (15) in liver-derived HepG2 cells, where PPARγ is highly expressed (16), revealed that PPARγ is indeed enriched at the *SETD6* promoter region in an open chromatin state (ATAC-seq), along with an enrichment for the histone marks H3K4me3 and H3K27ac in that genomic region (**Fig. 1B**). To determine whether PPARγ binds to putative PPRE within the promoter of *SETD6*, we performed a chromatin immunoprecipitation (ChIP) experiment to evaluate the occupancy of PPARγ in HepG2 cells in this genomic region. **Figure 1C** shows a significant enrichment of Flag-PPARγ at the *SETD6* promoter. Consistent with these results, over-expression of PPARγ resulted in a significant elevation in SETD6 mRNA expression level that was measured by qPCR **(Fig. 1D)**. To test the possibility that the observed occupancy on the *SETD6* promoter and its activation by PPARγ is direct, we cloned the full-length *SETD6* promoter upstream to a luciferase reporter gene (17), followed by transfection of this luciferase reporter construct and Flag-PPARγ in HepG2 cells. We found a significant elevation in *SETD6* promotor activity after over-expression of Flag-PPARγ **(Fig. 1E)**. Jointly, these results suggest that PPARγ binds to *SETD6* promoter and activates its transcription.

**Figure 1:**
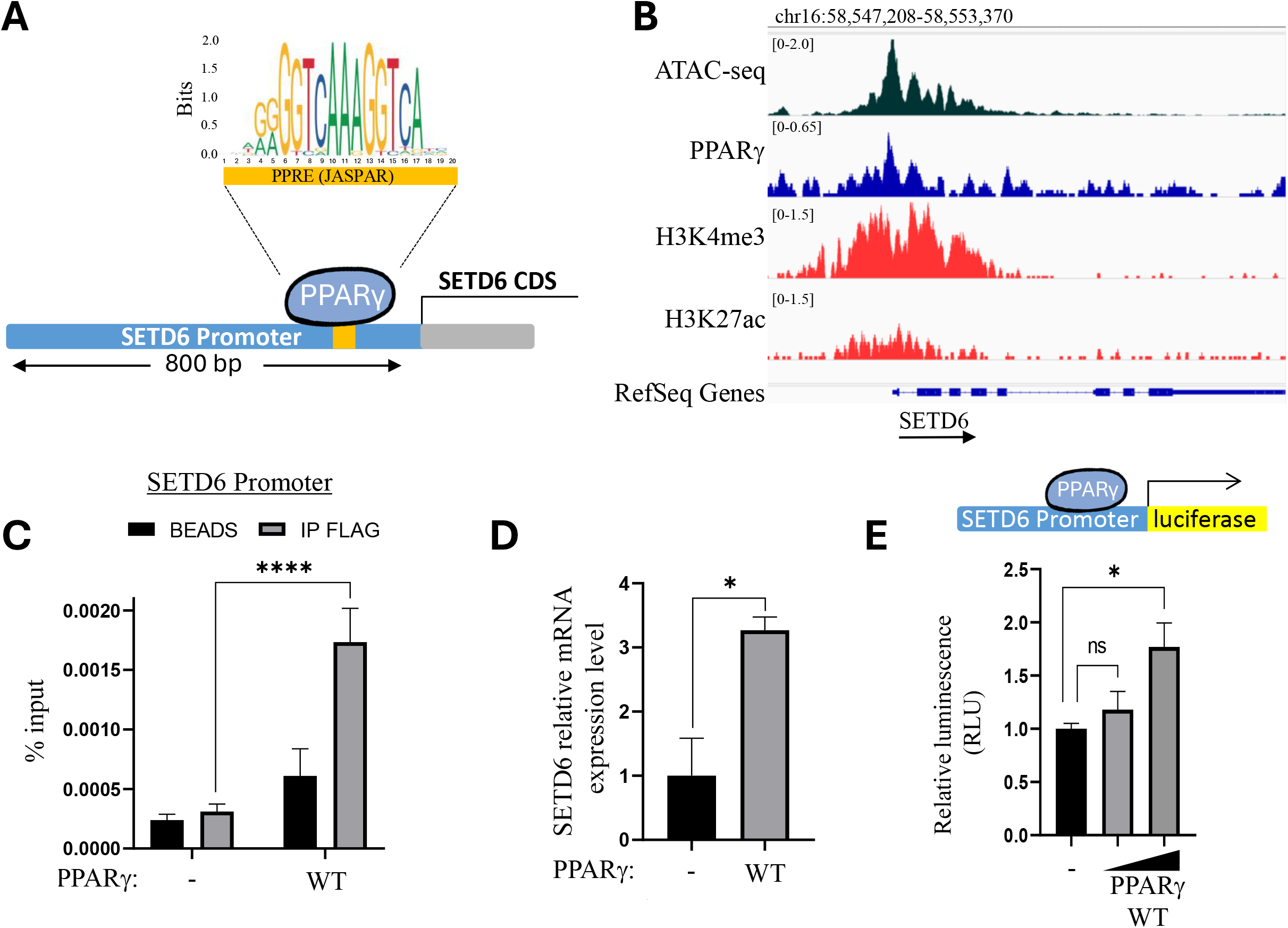
PPARγ binds SETD6 promoter and activates its expression. **(A)** Top: Sequence logo of the PPARγ response element (PPRE) from JASPAR. Bottom: Schematic representation of PPARγ binding to a predicted PPRE site within the SETD6 promoter **(B)** Capture of a genome browser showing the enrichment of PPARγ at the SETD6 promoter in HepG2 cells with open chromatin state represented by H3K4me3, H3K27ac, and ATAC-seq tracks. **(C)** ChIP assay with Flag-PPARγ antibody or beads as negative control in HepG2 cells followed by qPCR with primers flanking the predicted binding site at the SETD6 promoter. Graphs show % input of the quantified DNA. **(D)** RNA was extracted from HepG2 cells transfected with control or Flag-PPARγ WT. Transcript levels of SETD6 were determined by qPCR. Error bars are SEM. Statistical analysis was performed for three experimental repeats using one-way ANOVA (*p < 0.05, ****p < 0.0001). **(E)**- dual-luciferase assay in HepG2 cell transfect with an increasing amount of Flag-PPARγ. About 24 h post-transfection, the whole cell lysates were subjected to dual-luciferase assay (Promega). Relative luminescence was calculated after normalization of the firefly luciferase signal over Renilla luciferase control. Error bars are SD. Statistical analysis was performed for three experimental repeats. *p ≤ 0.03

### SETD6 binds and methylates PPARγ at K170 *in-vitro* and in cells

We have previously shown that SETD6 methylation of the transcription factor E2F1 positively regulates the expression level of SETD6 (17). We hypothesized that a similar mechanism occurs also with SETD6 and PPARγ. To address this hypothesis, we first tested if there is a direct physical interaction between SETD6 and PPARγ. An *in-vitro* ELISA experiment using purified recombinant proteins confirmed a direct SETD6-PPARγ interaction. MBP-RelA served as positive control (18,19) and BSA and PBS were used as negative controls (**Fig. 2A**). To determine whether SETD6 and PPARγ interact, we first examined their association under overexpression conditions. Co-expression of HA-SETD6 and Flag-PPARγ followed by FLAG immunoprecipitation from chromatin fractions, demonstrated an interaction between the two proteins (**Fig. S1**). To validate this interaction under endogenous expression level conditions, we next performed co-immunoprecipitation from RIPA lysates of HepG2 cells. Immunoprecipitation of endogenous PPARγ enriched endogenous SETD6, demonstrating an endogenous interaction between the two proteins (**Fig. 2B**). We next examined whether such interaction at endogenous expression levels also occurs on chromatin. Reciprocal co-immunoprecipitation experiments using chromatin-enriched fractions showed that endogenous SETD6 co-immunoprecipitated with endogenous PPARγ, confirming that the two proteins interact on chromatin (**Fig. 2C**). Finally, a proximity ligation assay (PLA) further validated the interaction in intact cells within the nuclei **(Fig. 2D)**. Notably, the catalytic inactive SETD6 mutant (Y285A) exhibited comparable proximity to PPARγ, indicating that SETD6 catalytic activity is not required for the physical association between the two proteins. Consistent with these findings, structural modeling of the SETD6–PPARγ complex predicted no major alterations in the overall interaction interface following substitution of Y285 with alanine **(Fig. S2)**, supporting a model in which SETD6 binding to PPARγ is independent of its catalytic activity.

**Figure 2:**
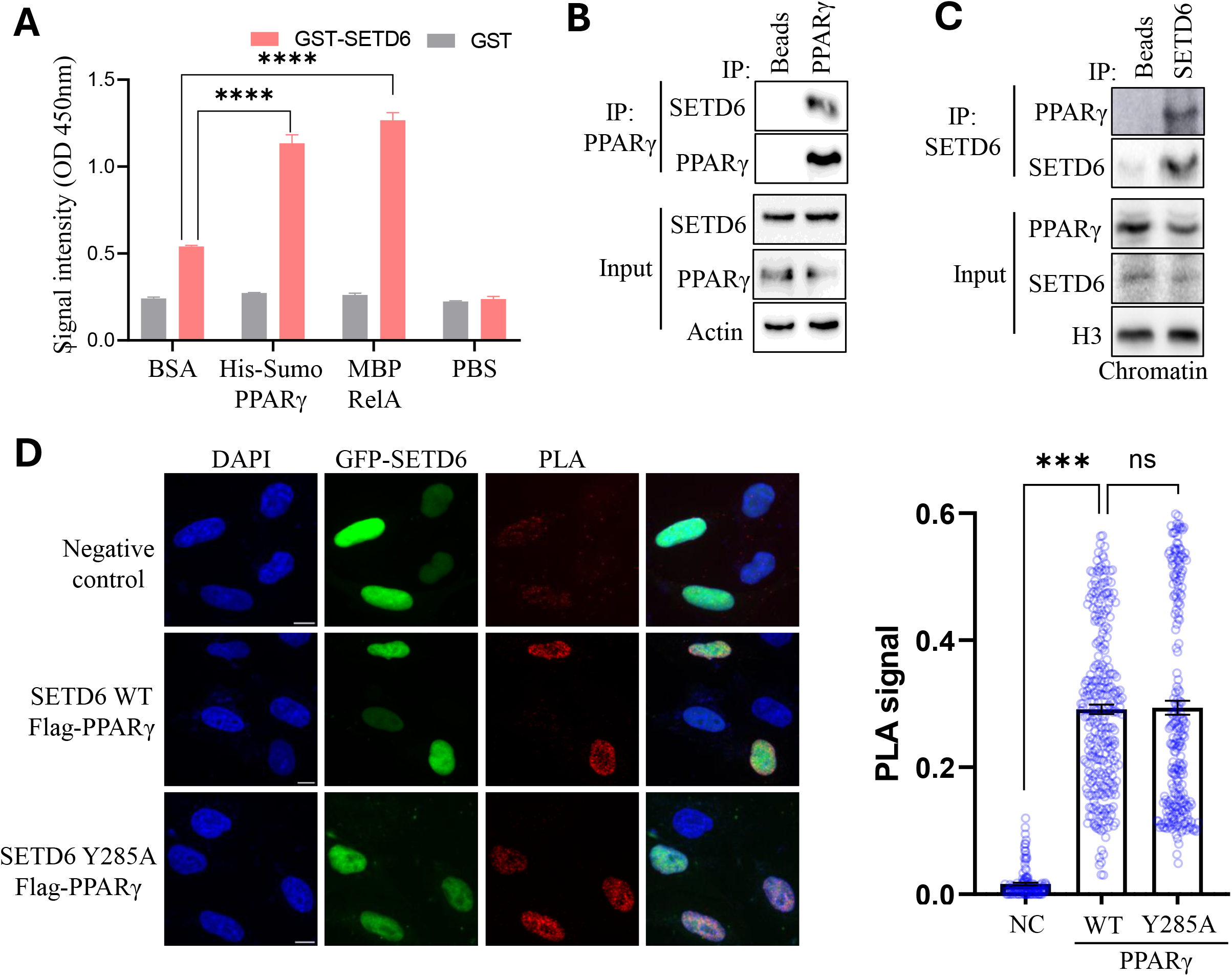
Physical interaction between PPARγ and SETD6 in-vitro and in cells. **(A)** ELISA-based analysis of the interaction between recombinant GST-SETD6 and the indicated recombinant proteins. ****p < 0.0001. **(B)** Endogenous PPARγ was immunoprecipitated from RIPA lysates of HepG2 cells followed by western blot analysis using the indicated antibodies. **(C)** Endogenous SETD6 was immunoprecipitated using chromatin fraction isolated from HepG2 followed by western blot analysis using the indicated antibodies. **(C)** Endogenous SETD6 was immunoprecipitated from chromatin isolated from HepG2 cells followed by western blot with the indicated antibodies. **(D)** Left, representative images of proximity ligation assays (PLA) between Flag-PPARγ and GFP-SETD6 WT or the catalytic inactive mutant GFP-SETD6 Y285A in HeLa cells. The negative control (NC) was performed in the absence of one primary antibody. Red dots indicate positive PLA signals. Scale bar = 10 μm. Right – Quantification of PLA signals for each sample. Statistical analysis was performed using Student’s t-test (***p < 0.001).

Given the physical interaction between SETD6 and PPARγ *in-vitro* and in cells, we hypothesized that PPARγ is methylated by SETD6. To address this hypothesis, recombinant PPARγ was subjected to an *in-vitro* methylation reaction using His-SETD6 and ^3^H -labeled SAM as the methyl donor. The results show that SETD6 methylates PPARγ (**Fig. 3A**). The methylation signal observed for SETD6 corresponds to its auto-methylation activity (20). To map the methylation site, *in vitro* methylation reactions were performed using non-radioactive SAM followed by mass spectrometry. The data obtained suggested, that PPARγ K170 mono-methylation (located in the DNA binding domain (DBD)) was the only methylation site identified after incubation with SETD6 (Spectra is shown in **Fig. S3**), and a schematic illustration of the DBD of PPARγ and the estimated location of K170 is shown in **Fig. 3A**, **top panel**). To validate this methylation site, we performed an *in-vitro* methylation reaction using a PPARγ with a K170R mutation. SETD6 failed to methylate the mutant PPARγ (**Fig. 3A**). Therefore, K170 is the major methylation site of PPARγ by SETD6. Immunoprecipitation with a pan-methyl antibody confirmed that endogenous PPARγ is methylated in HepG2 cells on chromatin (**Fig. 3B**). To test if this methylation is SETD6-dependent, we compared the methylation of Flag- PPARγ in CRISPR control (CT) and SETD6 knockout (KO) cells. The result revealed that Flag-PPARγ is methylated in cells in a SETD6-dependent manner (**Fig. 3C**). We next generated a site-specific antibody to detect PPARγ K170me1. Using a dot blot assay, we validated that the PPARγ K170me1 antibody specifically recognizes a PPARγ peptide mono-methylated at K170, but not the unmodified peptide (**Fig. 3D**). To confirm that PPARγ is methylated in cells on K170 by SETD6, we immunoprecipitated endogenous PPARγ from HepG2 cells in CRISPR CT and SETD6 KO cells, followed by WB with the PPARγ K170me1 antibody. A significant decrease in PPARγ methylation was observed in the SETD6 KO cells (**Fig. 3E**). Taken together, these results revealed that SETD6 binds and methylates PPARγ on K170 at chromatin *in vitro* and in cells.

**Figure 3:**
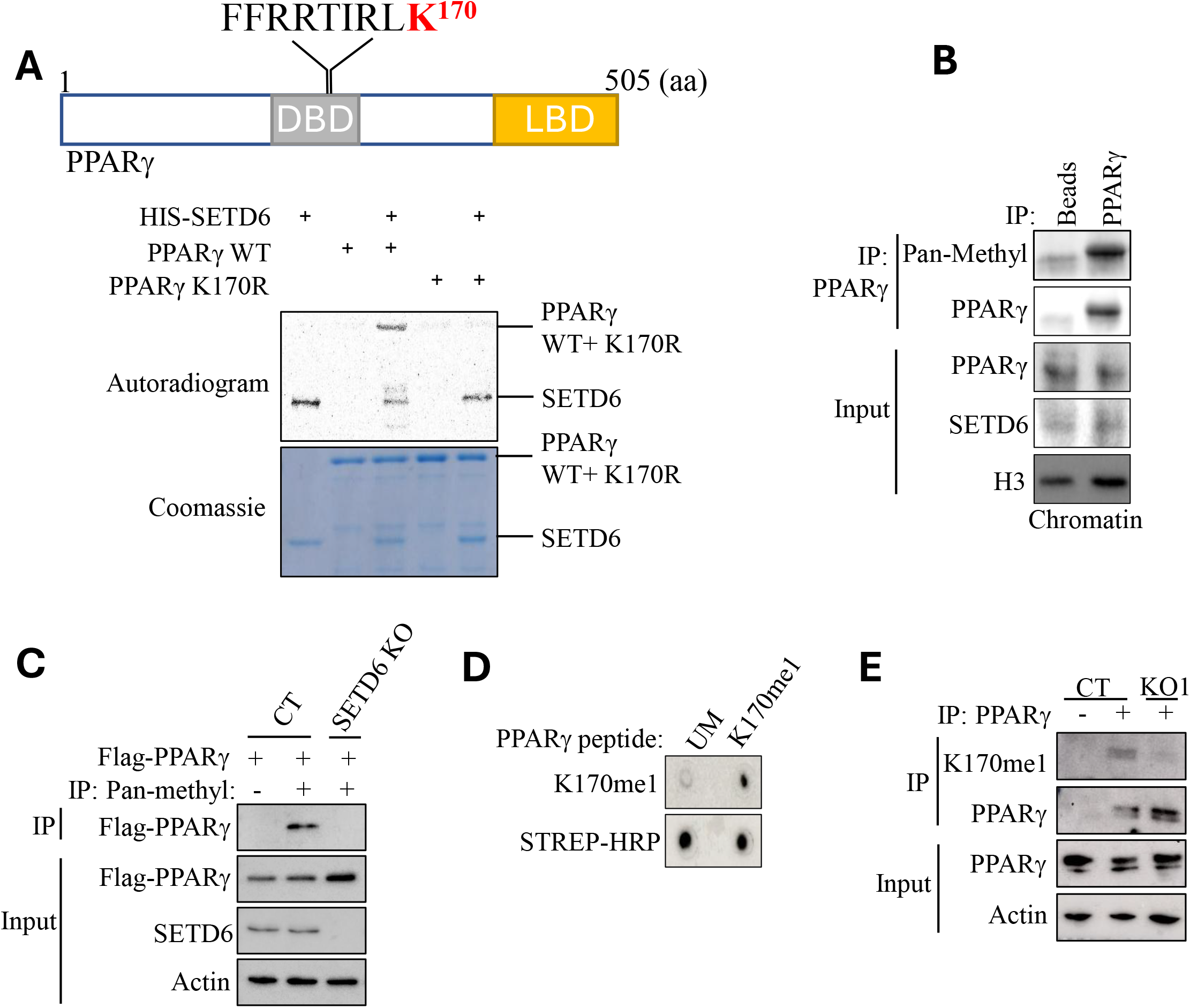
SETD6 methylates PPARγ at K170 *in-vitro* and in cells. **(A)** *In vitro* methylation assay in the presence of ^3^H-labeled SAM and the indicated purified proteins. Coomassie stain of the recombinant proteins used in the reactions is shown at the bottom. Schematic representation of PPARγ domain structure. The methylated residue (K170) identified by mass spectrometry is shown in red. DBD- DNA binding domain; LBD- ligand binding domain. **(B)** Endogenous PPARγ was immunoprecipitated from HepG2 cells followed by WB with the indicated antibodies **(C)** Flag-PPARγ was over-expressed followed by immunoprecipitation using pan-methyl antibody in control (CT) and SETD6 KO HeLa cells followed by western blot with indicated antibodies. **(D)** 0.25 ug of PPARγ peptides (un-modified and K170me1) were spotted on a nitrocellulose membrane followed by incubation with anti- PPARγ K170me1 antibody or Streptavidin-HRP. **(E)** Endogenous PPARγ was immunoprecipitated from control and KO SETD6 HepG2 cells followed by western blot with the indicated antibodies.

### PPARγ methylation at K170 regulates SETD6 promoter activation

Having demonstrated that PPARγ binds to the *SETD6* promoter and activates its transcription, we hypothesized that a molecular feedback mechanism may exist whereby SETD6-mediated K170 methylation of PPARγ affects SETD6 transcription. To test this hypothesis, we performed a luciferase reporter-based assay to determine *SETD6* promoter activity in control and SETD6 KO cells in the presence of over-expressed Flag-PPARγ. As shown in **Figure 4A**, knockout of SETD6 significantly decreased SETD6’s promotor activity. To complement these experiments, we measured SETD6 mRNA levels by qPCR in HepG2 cells stably expressing either Flag-PPARγ WT or the Flag-PPARγ K170R mutant. A significant elevation in SETD6 mRNA expression level was observed in cells overexpressing WT PPARγ compared with the PPARγ K170R mutant (**Fig. 4B)**. To determine whether the occupancy of PPARγ on the *SETD6* promotor is SETD6-dependent, we performed ChIP experiments of endogenous PPARγ in HepG2 CRISPR CT and SETD6 KO cells (**Fig. 4C).** SETD6 KO led to a significant reduction in the occupancy of PPARγ on the SETD6 promoter. To test if the occupancy of PPARγ on the *SETD6* promoter is K170me1-dependent, a ChIP experiment was performed in cells stably expressing Flag-PPARγ WT and Flag-PPARγ K170R mutant (**Fig. 4D)**. The enrichment of Flag-PPARγ was significantly higher in the cells expressing PPARγ WT compared to the K170R mutant. Taken together, the results suggest that PPARγ binds to the *SETD6* promoter and activates SETD6 mRNA expression in a SETD6-mediated, K170me1-dependent manner, thereby demonstrating a positive feedback loop.

**Figure 4:**
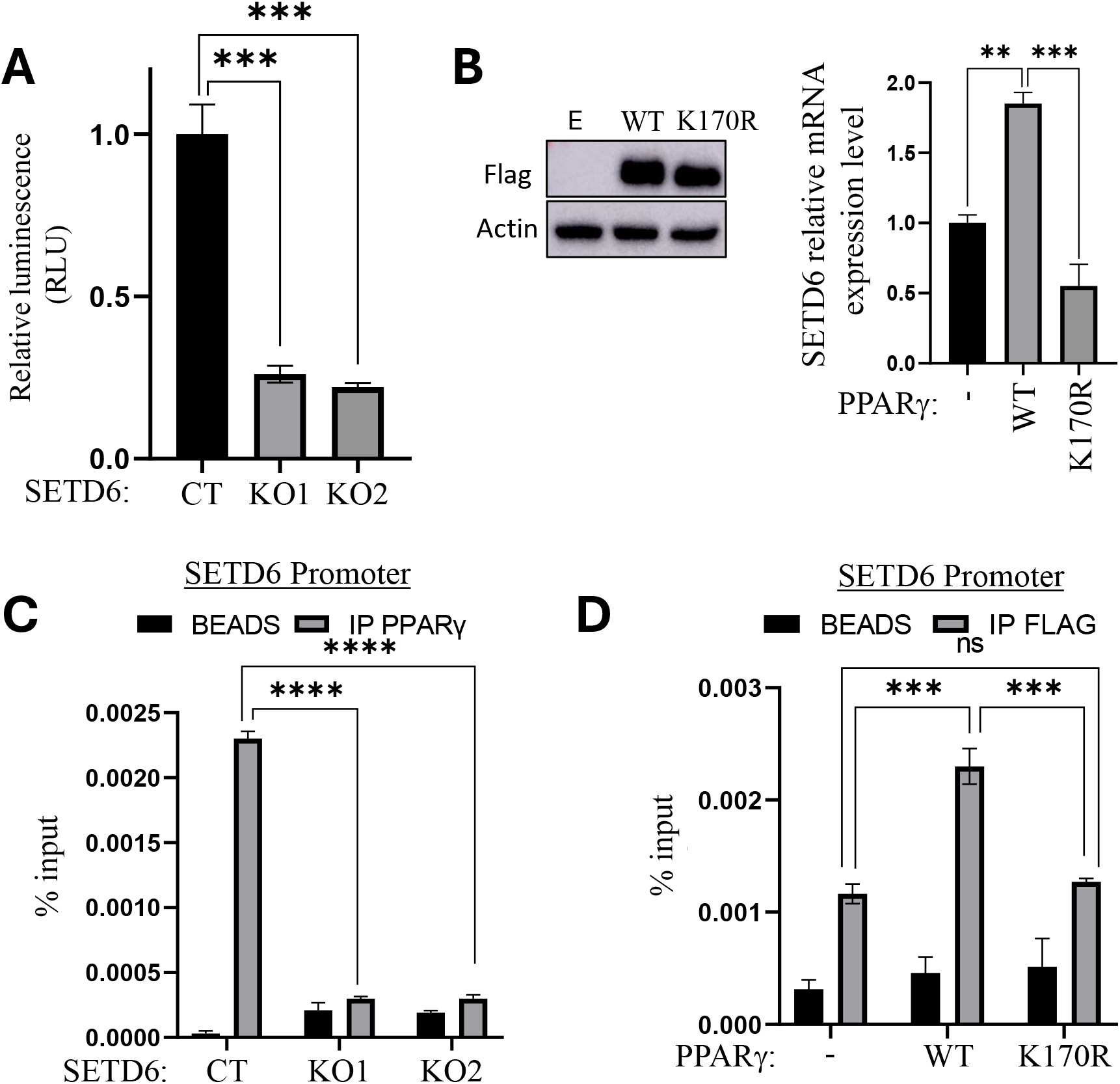
PPARγ binds and activates SETD6 expression in a K170 methylation dependent manner. **(A)** - dual-luciferase assay in Control and SETD6 KO HepG2 cells. 24 h post-transfection, the whole cell lysates were subjected to dual-luciferase assay (Promega). Relative luminescence was calculated after normalization of the firefly luciferase signal over Renilla luciferase control. Error bars are SD. Statistical analysis was performed for three experimental repeats. *\*\*\*p* < 0.001. **(B)** Left- WB analysis with the indicated antibodies for stable cells expressing Flag-WT or Flag-K170R PPARγ. Right- Transcript levels of the SETD6 were determined by qPCR of stably expressing HepG2 cells- Empty, Flag-PPARγ WT, and Flag-PPARγ K170R mutant. mRNA levels were normalized to GAPDH and then to Empty. **(C)** Chromatin immunoprecipitation (ChIP) assay. The chromatin fraction of HepG2 SETD6 CRISPR CT, KO1, and KO2 cells were immunoprecipitated with magnetic beads conjugated with anti-PPARγ antibody. The bound DNA was purified and amplified by qPCR. **(D)** Same as C for cells stably expression of Empty, Flag-PPARγ WT, and Flag-PPARγ K170R mutant. Graphs for C and D show the percent input of the quantified DNA. Two-way ANOVA analysis was performed; error bars are S.E.M. *\*p* < 0.05; *\*\*p* < 0.01; *\*\*\*p* < 0.001; *\*\*\*\*p < 0.0001*.

### PPARγ K170 methylation by SETD6 regulates lipid droplet formation

To assess the global transcriptomic impact of SETD6 on human hepatocellular carcinoma cells in an unbiased manner, we performed RNA sequencing in control HepG2 cells (2 independent gRNAs) and SETD6 knock-out (KO) cells (derived from 3 independent gRNAs). The KO efficiency of SETD6 gRNAs in these cells is shown in **Fig. 5A**. RNA-sequencing analysis identified 302 differentially expressed genes in SETD6 KO HepG2 cells compared to control cells, including 166 downregulated and 136 upregulated genes (**Fig. 5B**). To better define the biological pathways associated with each gene set, KEGG pathway enrichment analysis was performed separately for the downregulated **(Fig. 5C**) and upregulated genes **(Fig. 5D**). Genes downregulated in SETD6 KO cells were significantly enriched for pathways associated with lipid metabolism, including the PPAR signaling pathway, glycerolipid metabolism, cholesterol metabolism, and fat digestion and absorption. Additional enriched pathways included TNF signaling, focal adhesion, and pluripotency-related pathways, which are consistent with previously reported roles of SETD6 in transcriptional regulation and cancer-associated signaling (12) (**Fig. 5C).** In contrast, genes upregulated in SETD6 KO cells were enriched for pathways associated with endoplasmic reticulum function, oxidative phosphorylation, lysosomal pathways, and glycerophospholipid **(Fig. 5D**). Together, these findings further support a functional role for SETD6 in regulating PPARγ-dependent transcriptional programs and lipid metabolic pathways in hepatocellular carcinoma cells. To phenotypically validate whether SETD6 has a role in fat accumulation, we established a live cell imaging system that allows us to quantify and study the kinetics of lipid droplet accumulation in live cells (**Fig. 5E)**. In this system, we treated HepG2 with oleic acid (OA), as the primary fatty acid source (21) followed by staining with BODIPY 493/503, a green-fluorescent dye that stains neutral lipids (22). Hoechst dye (23) was used to identify the nuclei and quantify the number of cells, enabling a calculation of mean green fluorescence over time. First, we conducted a 20-hour calibration experiment to determine the appropriate OA concentration (**Fig. S4**). We incubated HepG2 cells with three different concentrations of OA; 300, 600 and 900 μM, and with DMSO serving as a vehicle control. OA accumulation in the cells treated with 300 μM and 600 μM reached a plateau over time. The DMSO-treated cells exhibited a decline in LD accumulation, indicating that the BODIPY dye is specific to neutral lipid staining. We chose to work with the 600uM concentration of OA since it yielded optimal staining conditions and did not affect cell morphology. We next asked if depletion of SETD6 affects the formation of lipid droplets in cells? To this end, HepG2 CT and KO SETD6 cells were treated with 600 uM of OA over 24 hours, and the mean GFP signal was measured. The result shows that the CT cells accumulated significantly more lipid droplets over time compared to SETD6 KO1 and KO2 cells (**Fig. 5F**). Representative images of HepG2 CRISPR CT, KO1, and KO2 cells at a 24-hour time point are presented (**Fig. 5F, top panel**). To confirm that the lipid droplets accumulation is dependent on SETD6, we performed a rescue experiment wherein SETD6 KO cells were transfected with HA-SETD6 plasmid **(Fig. 5H**; SETD6 expression is confirmed by western blot in **Fig. 5G**). The results showed that exogenous HA-SETD6 expression reversed the effect of SETD6 knockout. These findings suggest that SETD6 positively regulates lipid droplet formation.

**Figure 5:**
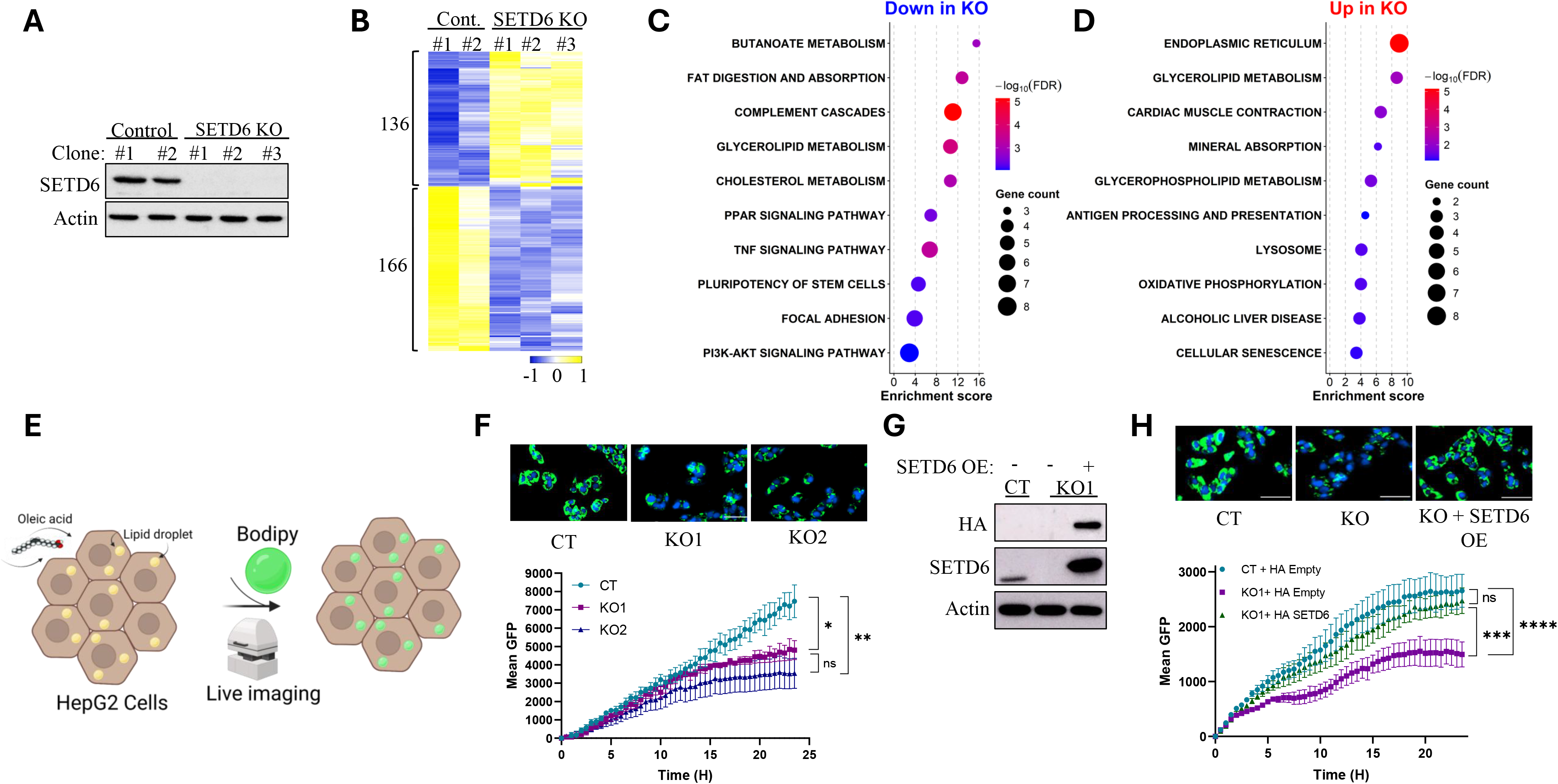
SETD6 positively regulates lipid droplets formation: **(A)** WB analysis with the indicated antibodies for HepG2 control (2 clones) and SETD6 KO cells (3 clones) **(B)** Heatmap showing upregulated and downregulated genes identified by RNA-sequencing analysis of two SETD6 control (CT) and three SETD6 KO HepG2 independent clones. Gene expression values are displayed as normalized Z-scores. Yellow and blue colors represent relatively high and low expression levels, respectively. **(C+D)** KEGG pathway enrichment analysis of differentially expressed genes identified by RNA-sequencing analysis of SETD6 control (CT) and SETD6 KO HepG2 cells. **(C)** pathways enriched among genes downregulated in SETD6 KO cells. **(D)** pathways enriched among genes upregulated in SETD6 KO cells. Circle size represents the number of differentially expressed genes associated with each pathway, and color intensity indicates statistical significance expressed as −log10(FDR). The x-axis represents the enrichment score. **(E)** Illustration of the system to monitor lipid droplets accumulation over time: HepG2 cells are treated with oleic acid (OA) and stained with Hoechst (nucleus) and BODIPY(neutral lipids), followed by live cell imaging. **(F)** Top-Representative images of HepG2 CRISPR CT (control), KO1, and KO2 challenged with 300 mM of OA (20 H time point) and stained with Hoechst (nucleus) and BODIPY (neutral lipids). Bottom- Mean green fluorescence was calculated as fluorescence signal divided by cell count. Data is analyzed from three beacons per well in three wells. Statistical analysis was performed using one-way ANOVA. ns: non-significant; *p < 0.05; **p < 0.01. **(G)** WB analysis for CT and SETD6 KO with or without over-expression of HA-SETD6. **(H)** Top- Representative images of HepG2 CRISPR CT (control) and SETD6 KO without or with over-expression of HA-SETD6, challenged with 300 mM of OA (20 H time point) and stained with Hoechst (nucleus) and BODIPY (neutral lipids). Bottom- Mean green fluorescence was calculated as fluorescence signal divided by cell count. Data is analyzed from three beacons per well in three wells. Statistical analysis was performed using one-way ANOVA. ns: non-significant; ***p < 0.001; ****p < 0.0001.

Given that SETD6 methylates PPARγ and positively regulates lipid droplet formation, we hypothesized that this phenotype is mediated through PPARγ K170 methylation and its transcriptional activity. To determine whether SETD6-mediated methylation of PPARγ at K170 regulates gene expression in HepG2 cells, we performed RNA-sequencing analysis using HepG2 cells stably expressing Flag-PPARγ WT or the Flag-PPARγ K170R mutant (**Fig. 6A**). We identified 1,122 differentially expressed genes, including 532 downregulated and 590 upregulated genes in the PPARγ K170R mutant cells compared to WT PPARγ-expressing cells.

**Figure 6:**
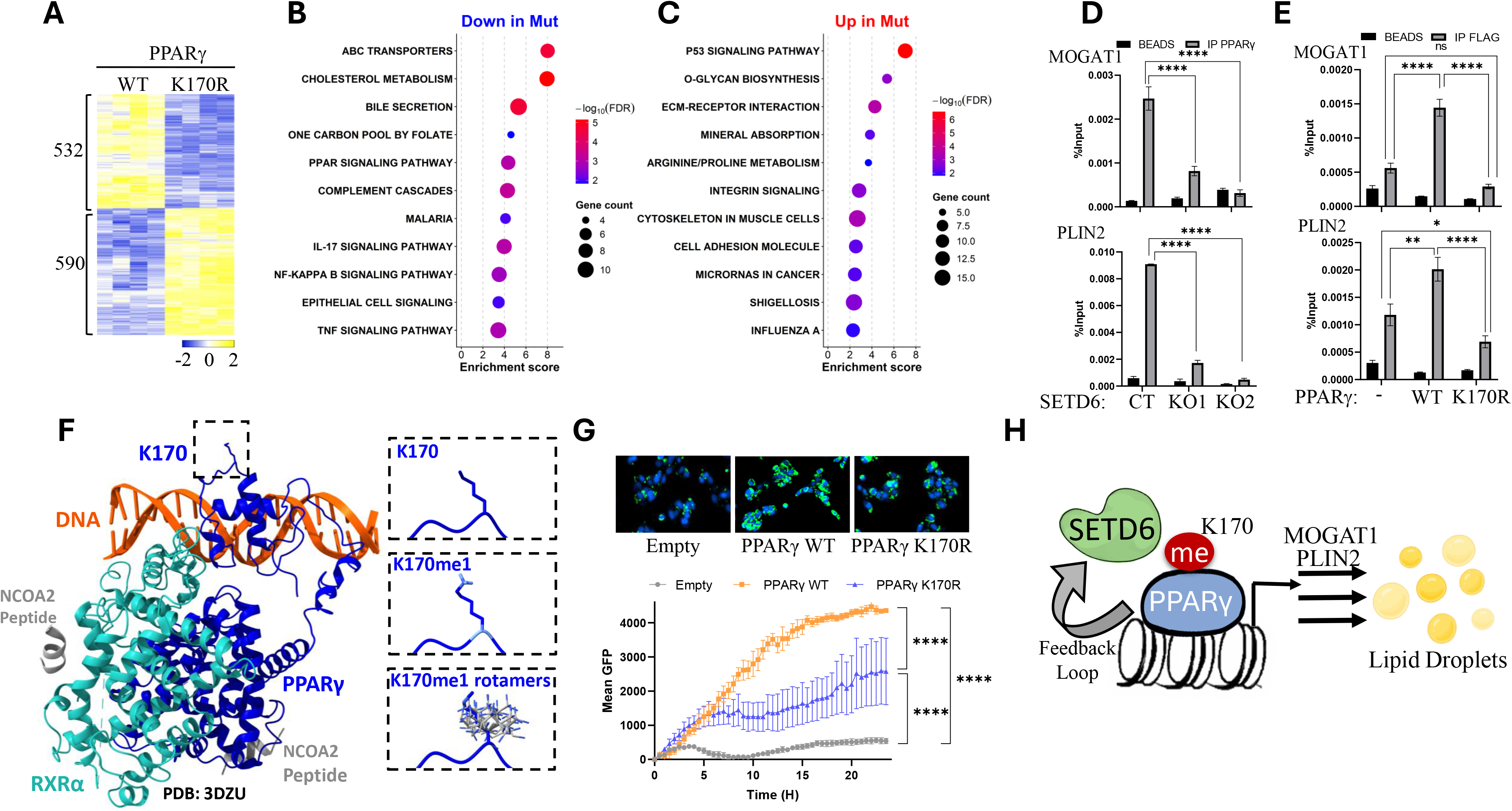
PPARγ methylation at K170 positively regulates gene expression and lipid droplets formation. **(A)** Heatmap showing upregulated and downregulated genes identified by RNA-sequencing analysis of HepG2 independent clones stably expressing PPARγ WT or the PPARγ K170R mutant. Gene expression values are displayed as normalized Z-scores. Yellow and blue colors represent relatively high and low expression levels, respectively. **(B+C)** KEGG pathway enrichment analysis of differentially expressed genes identified by RNA-sequencing analysis of HepG2 cells stably expressing PPARγ WT or the PPARγ K170R mutant. **(B)** pathways enriched among genes downregulated in the PPARγ K170R mutant cells. **(C)** pathways enriched among genes upregulated in the PPARγ K170R mutant cells. Circle size represents the number of differentially expressed genes associated with each pathway, and color intensity indicates statistical significance expressed as −log10(FDR). The x-axis represents the enrichment score. **(D)** ChIP analysis for HepG2 control and SETD6 KO cells (two clones) that were immunoprecipitated with PPARγ antibody. The bound DNA was purified and amplified by qPCR using specific primers to MOGAT1 and PLIN2 gene promoter regions. Graphs show the percent input of the quantified DNA. Two-way ANOVA analysis was performed; error bars are S.E.M. ****p < 0.0001. **(E)** Same as for **panel D** with HepG2 cells stably expressing Empty, Flag-PPARγ2 WT, and Flag-PPARγ2 K170R mutant that were immunoprecipitated with FLAG conjugated magnetic beads. error bars are S.E.M. ns: non-significant; *\*p* < 0.05; *\*\*p* < 0.01; *\*\*\*\*p < 0.0001*. **(F)** Structural model in cartoon representation of the PPARγ–RXRα heterodimer bound to DNA based on the published co-crystal structure (PDB: 3DZY). PPARγ is shown in blue, RXRα in cyan, DNA in orange, and NCOA2 peptides in gray. The boxed region indicates K170 within the PPARγ DNA-binding domain. Enlarged views in stick representation of the K170 side chain (top), the modeled K170me1 side chain (middle), and multiple modeled K170me1 rotamers (bottom) are shown. **(G)** Top- Representative images of HepG2 cells stably expressing Empty, PPARγ WT or PPARγ K170R mutant, challenged with 300 mM of OA (20 H time point) and stained with Hoechst (nucleus) and BODIPY (neutral lipids). Bottom- Mean green fluorescence was calculated as the fluorescence signal divided by cell count. Data is analyzed from three beacons per well in three wells. Statistical analysis was performed using one-way ANOVA. ****p < 0.0001. **(H)** A Schematic representation of our proposed working model.

To better characterize the biological pathways associated with PPARγ K170 methylation, KEGG pathway enrichment analysis was performed separately for the downregulated and upregulated gene sets (**Fig. 6B and 6C**). Genes downregulated in the PPARγ K170R mutant cells (**Fig. 6B)** were significantly enriched for pathways associated with lipid metabolism, including the PPAR signaling pathway, cholesterol metabolism, bile secretion, and ABC transporters. Additional enriched pathways included TNF signaling, NF-κB signaling, and IL-17 signaling pathways. In contrast, genes upregulated in the PPARγ K170R mutant cells (**Fig. 6C)** were enriched for pathways associated with p53 signaling, ECM-receptor interaction, integrin signaling, and cell adhesion-related pathways. These findings are consistent with the enriched pathways identified by RNA-sequencing analysis of control and SETD6 KO cells (**Figure 5C**), further supporting a functional role for SETD6-mediated PPARγ methylation in regulating lipid metabolic transcriptional programs. The RNA-seq results were further validated by qPCR (**Fig. S5**). Taken together, this data demonstrates that methylation of PPARγ at K170 positively regulates the expression of genes associated with lipid droplet formation.

The fact that K170 is in the DNA binding domain of PPARγ, may suggest that the methylation affects its association with target genes linked to lipid droplets formation. To test this hypothesis, we decided to focus on two PPARγ target genes, that are known to be involved in the LD formation process; MOGAT1 - a rate-limiting enzyme involved in incorporating fatty acids into triglycerides and characterizes the fatty-acid esterification step (24), and PLIN2 - a lipid droplet-associated protein involved in storage of neutral lipids within lipid droplets (25). ChIP experiments revealed that PPARγ is significantly more enriched at the MOGAT1 and PLIN2 loci in the SETD6 CT cell line compared to the SETD6 KO cells (**Fig. 6D**). Consistent with these results, using HepG2 stable cell lines we observed a greater enrichment of FLAG-PPARγ WT on both MOGAT1 and PLIN2 promoter regions in comparison to the FLAG-PPARγ K170R mutant (**Fig. 6E**). Because K170 resides within the DNA-binding domain of PPARγ, we next asked whether methylation at this residue could directly influence the interaction between PPARγ and DNA. Structural modeling based on published co-crystal structure of PPARγ bound to DNA (PDB: 3DZY) (26) demonstrates that K170me1 side chain is oriented away from the DNA interface and does not introduce steric clashes with DNA (**Figure 6F**). Furthermore, modeling of multiple K170me1 rotamers yielded similar results, suggesting that K170 methylation is unlikely to directly alter the DNA-binding interface through steric effects. Together with our ChIP-qPCR data, these findings support a model in which K170 methylation promotes PPARγ chromatin occupancy through indirect mechanisms, such as modulation of protein–protein interactions, cofactor recruitment, or local chromatin organization, rather than by directly affecting the DNA-binding interface.

Finally, to test whether lipid droplet formation is dependent on PPARγ methylation at K170, we subjected the cells to the live imaging system described earlier. Lipid droplets accumulation was significantly lower in cells expressing Flag-PPARγ K170R compared to Flag-PPARγ WT (**Fig. 6G**). These results strongly suggest that K170 PPARγ methylation positively mediates lipid droplet formation. Taken together, we propose a working model in which SETD6 methylates PPARγ at lysine 170 (K170), enhancing its transcriptional activity on target genes involved in lipid droplet formation, including *MOGAT1* and *PLIN2*. This activation promotes lipid droplet accumulation. In parallel, PPARγ K170 methylation drives SETD6 transcription, establishing a positive feedback loop that sustains the expression of LD-associated genes (**Fig. 6H**).

## Discussion

Previous studies have shown that the transcriptional activity of PPARγ is significantly regulated by various types of post-translational modifications, including phosphorylation, acetylation, ubiquitination, and sumoylation (27–31). Interestingly, a recent paper has shown that SMYD2-miediated mono methylation of PPARγ facilitates hypoxia-induced pulmonary hypertension and promotes mitophagy (32). However, the methylation site has not been identified. Here, we identify a regulatory feed-forward circuit between PPARγ and the lysine methyltransferase SETD6, establishing SETD6 as a previously unrecognized enzymatic regulator of PPARγ transcriptional activity. We show that SETD6 mono-methylates PPARγ at K170, a lysine residue that resides within its DNA-binding domain, thereby promoting PPARγ chromatin occupancy, the expression of genes involved in lipid metabolism, and lipid droplet biogenesis in HepG2 cells. Consistently, analysis of previously generated PPARγ ChIP-seq experiments (15) revealed PPARγ occupancy at the SETD6 promoter. We demonstrate that K170 methylation enhances PPARγ occupancy at this promoter, resulting in increased SETD6 transcription and mRNA expression. Together, these findings establish a positive feed-forward regulatory loop in which SETD6 activates PPARγ, which in turn transcriptionally upregulates SETD6 expression. Our results do not only suggest that PPARγ positively regulates the expression of SETD6 but also provide evidence that it occurs in a SETD6- and a PPARγ methylation-dependent manner. Our previous work demonstrated that SETD6 methylates E2F1 at K117 (17), and that this modification promotes SETD6 gene expression, establishing a positive regulatory circuit reminiscent of the one we describe herein, involving PPARγ. Thus, SETD6 appears to participate in functional transcriptional feed-forward loops involving distinct transcription factors. Although the upstream pathways controlling SETD6 expression remain largely unknown, the recurring ability of SETD6 to regulate transcription factors that, in turn, promote its own expression suggests that tight control of SETD6 abundance and enzymatic activity is an important mechanism for coordinating context-dependent transcriptional programs.

Our RNA sequencing analysis of stably expressing HepG2 cells with wild-type PPARγ and the PPARγ K170R mutant provided insight into the functional consequences of SETD6-mediated methylation on PPARγ in various biological processes. Specifically, we observed that methylation at lysine 170 (K170) positively regulates the expression of genes associated with lipid metabolism, particularly those involved in lipid droplet formation. These findings highlight the critical role of PPARγ methylation in modulating metabolic pathways linked to lipid storage and processing. Given that the K170 methylation site is located within the DNA-binding domain (DBD) of PPARγ, specifically between its two zinc-finger motifs, we investigated how this post-translational modification affects the occupancy of PPARγ at its target gene promotors. Our analysis focused on its binding to peroxisome proliferator response elements (PPREs) present in the promoters of key target genes, including MOGAT1 and PLIN2 (33,34).

Consistent results were observed in both SETD6 CRISPR knockout cells and overexpression models. PPARγ exhibited significantly greater enrichment in the promoter regions of control SETD6 CRISPR cells (in comparison to SETD6 KO) and cells stably expressing Flag-PPARγ WT cells (compared to Flag-PPAR K170R mutant). These findings demonstrate that the methylation of PPARγ at K170 by SETD6 facilitates its recruitment to promoter region of its target genes. The increased transcriptional activity of MOGAT1 and PLIN2 likely drives their protein expression, with the functional results being enhanced lipid accumulation. A broader investigation into additional PPARγ target genes that may be affected by SETD6 expression and K170 methylation is required to further understand whether our finding reflect a more generalized phenomenon

Lysine K170 on PPARγ is conserved among the other PPAR family member, PPARβ/δ but not in PPARα. These findings suggest that the other PPAR family members, which have other physiological roles in different tissues (35–37), can be similarly regulated by SETD6 and methylation. Future experiments will determine whether the newly identified methylation event is redundant or is PPARγ-specific.

Although our structural modeling suggests that K170 methylation is unlikely to directly alter the DNA-binding interface through steric effects **(Figure 6F)**, it may influence local conformational dynamics of the DNA-binding domain or modulate interactions with transcriptional cofactors and chromatin-associated proteins. Given that K170 is located between the two zinc-finger motifs of the DNA-binding domain, this modification may contribute to selective chromatin occupancy, promoter recognition, or stabilization of transcriptional complexes at PPARγ target genes. Future biochemical, mechanistic, and structural studies will be required to distinguish between these possibilities.

Based on the PhosphoSite database, K170 of PPARγ has not been reported to undergo additional post-translational modifications, such as acetylation or ubiquitination. Interestingly, however, the neighboring residue threonine 166 (T166), located within the DNA-binding domain, is phosphorylated by PKCα and has been shown to regulate PPARγ transcriptional activity and lipid metabolic programs in adipocytes (38). To further explore this possibility, we performed structural modeling based on the published PPARγ–DNA co-crystal structure (PDB: 3DZY). The model predicts that the K170me1 side chain is oriented away from the DNA interface and does not introduce steric clashes with DNA, suggesting that K170 methylation is unlikely to directly alter the DNA-binding interface. In contrast, phosphorylation of T166 is predicted to introduce multiple intramolecular steric clashes within the DNA-binding domain, which may alter its local conformation or dynamics and thereby indirectly influence DNA binding or transcriptional activity **(Figure S6)**. Together with our ChIP-qPCR data demonstrating reduced promoter occupancy of the PPARγ K170R mutant, these observations support a model in which K170 methylation primarily regulates PPARγ chromatin occupancy through indirect mechanisms, such as modulation of protein–protein interactions, cofactor recruitment, or local chromatin organization, rather than through direct steric effects on the DNA-binding interface. While these observations remain speculative and require further experimental validation, they suggest that phosphorylation and methylation within the DBD may coordinately regulate PPARγ chromatin occupancy and transcriptional selectivity.

Accumulating evidence suggests that SETD6 functions as a broader regulator than previous appreciated of transcription factor activity through lysine methylation. Previous studies demonstrated that SETD6-mediated methylation modulates the activity of multiple chromatin-associated factors, including RelA, E2F1, TWIST1, and BRD4 (12,17,18,39–41), thereby influencing chromatin organization, cofactor recruitment, and transcriptional selectivity. Together with the findings presented here, these observations raise the possibility that SETD6 acts as a context-dependent epigenetic signaling regulator that integrates metabolic, inflammatory, and oncogenic cues through selective methylation of transcription factors. In the context of PPARγ, it is possible that conditions associated with elevated fatty acid flux, steatosis, or metabolic stress promote the functional significance of PPARγ K170 methylation, potentially by stabilizing chromatin occupancy or facilitating recruitment of additional transcriptional cofactors. Further studies will be required to define the upstream signals and physiological conditions that regulate SETD6 catalytic activity toward distinct substrates.

Nonalcoholic fatty liver disease (NAFLD), recently also termed metabolic dysfunction-associated fatty liver disease (MAFLD), is the most common chronic liver disorder worldwide, affecting approximately 25% of the global population (42). It represents a spectrum of liver diseases characterized by abnormal accumulation of triglycerides (TGs) and lipid droplets within hepatocytes, and is clinically defined by lipids accounting for more than 5% of the liver weight (43). Lipid droplets are the primary intracellular storage organelles for neutral lipids and are predominantly composed of triacylglycerols (TAGs) and steryl esters (STEs) surrounded by a phospholipid monolayer (44–46). A central mechanism contributing to hepatic steatosis occurs when adipose tissue exceeds its lipid storage capacity, leading to elevated basal adipocyte lipolysis and increased release of free fatty acids into circulation. This lipid overflow promotes excessive fatty acid uptake by hepatocytes and the formation of lipid droplets within the hepatocytes. While steatosis initially develops as a metabolic adaptation to nutrient excess, persistent lipid accumulation can trigger oxidative stress, inflammation, and cellular dysfunction. In a substantial subset of patients, MAFLD progresses to steatohepatitis (MASH), fibrosis, cirrhosis, greatly increasing the risk for hepatocellular carcinoma (47–49). These emphasize the need to better understand the molecular pathways regulating hepatic lipid accumulation and metabolic stress responses.

Future studies should therefore involve physiologically relevant in-vivo models used to investigate MAFLD progression and metabolic liver disease. Such approaches will allow us to evaluate the role of SETD6-mediated PPARγ K170 methylation under diet-induced metabolic stress conditions and to further validate the mechanistic model proposed in this study. In particular, mice expressing either wild-type PPARγ or the K170R mutant, whole body or in a tissue-specific manner, will enable assessment of the whole-body impact of SETD6-dependent methylation of PPARγ via genomic programs, lipid droplet accumulation, steatosis progression, and inflammatory responses.

In summary, this study highlights the pivotal cellular role of SETD6 and its lysine methylation activity via a central regulator of cellular metabolism - PPARγ. Our findings uncover a previously unrecognized function of SETD6 in promoting lipid droplets accumulation, mediated through its influence on the DNA-binding capacity and transcriptional regulation of PPARγ target genes in a methylation dependent manner. Integrating the data obtained in this study will pave new research directions to gain more insight into SETD6’s role in metabolic diseases through the methylation of PPARγ, with potential therapeutic applications.

## Materials and Methods

### Cell lines treatment

HepG2 (Hepatocellular Carcinoma) cells were maintained in Eagle’s Minimum Essential Medium (EMEM) (BI, 01-025-1A) with 10% fetal bovine serum (FBS) (Gibco, 10270106), 2 mg/ml L-glutamine (Sigma, G7513) and 1mM sodium pyruvate (BI, 03-042-1B). Human embryonic kidney cells (HEK-293T) and HeLa cells were maintained in Dulbecco’s modified Eagle’s medium (DMEM) (Sigma, D5671) with 10% fetal bovine serum (Gibco, 10270106), 2 mg/ml L-glutamine (Sigma, G7513),1% penicillin-streptomycin (Sigma, P0781), and non-essential amino acids (Sigma, M7145). Cells were cultured at 37°C in a humidified incubator with 5% CO2.

### Plasmids

Mouse PPARγ2 (mPPARγ2) coding sequence was excised from plasmids that were kindly provided by Yosef Shaul’s lab from Weizmann Institute of Science. In addition, the human PPARγ2 (hPPARγ2) coding sequence was amplified from a sample of cDNA sequence from human visceral adipose tissue that was kindly provided by Prof. Assaf Rudich’s lab. mPPARγ2 and hPPARγ2 sequences were amplified by using PCR with compatible primers as indicated in Table 1. The amplicons were digested with AscI and PacI restriction enzymes and mPPARγ2 subcloned into a pET-Sumo plasmid for protein purification. mPPARγ2 and hPPARγ2 subcloned into pcDNA3.1 3xFlag. To generate mPPARγ2 and hPPARγ2 mutants, site-directed mutagenesis on a PPARγ2 wild-type vector was performed for each plasmid using other primers indicated in Table 1. PPARγ2 K170R mutants were also cloned into pcDNA3.1 3xFlag, and mPPARγ2 K170R mutant was cloned into a pET-Sumo plasmid for protein purification. PCR reactions were accomplished using the KAPA HiFi HotStrart ReadyMix (KAPA Biosystems), and cloned plasmids were confirmed by sequencing. Cloning pET-Duet and pcDNA3.1 hSETD6 (Human) plasmid containing His and HA tags described in (50), respectively. *SETD6* promoter sequence was amplified by PCR using primers indicated in table 1, and subcloned into pGL3 plasmid.

**Table 1.**
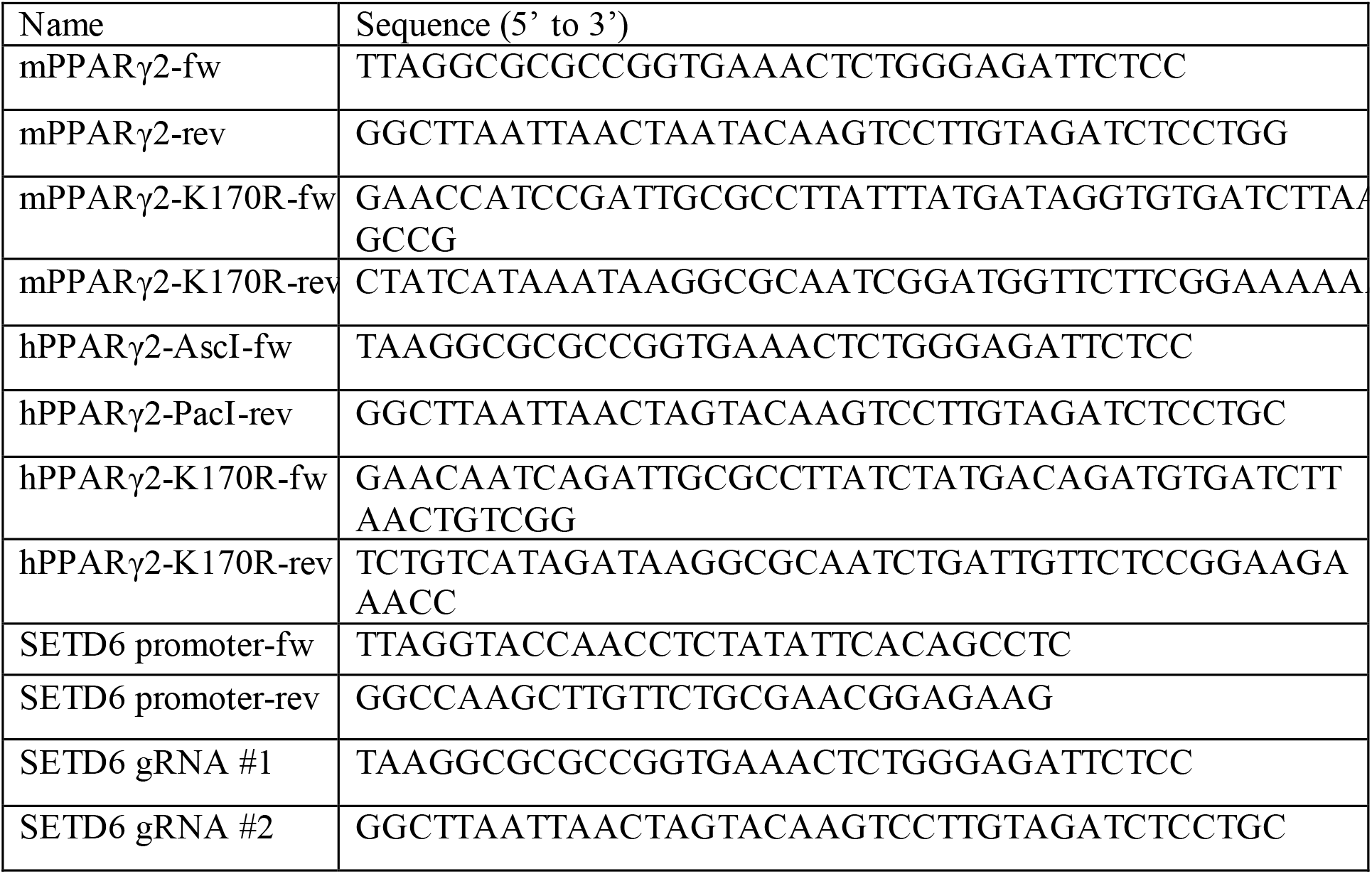
Primers for cloning and mutagenesis.

**Table 2.**
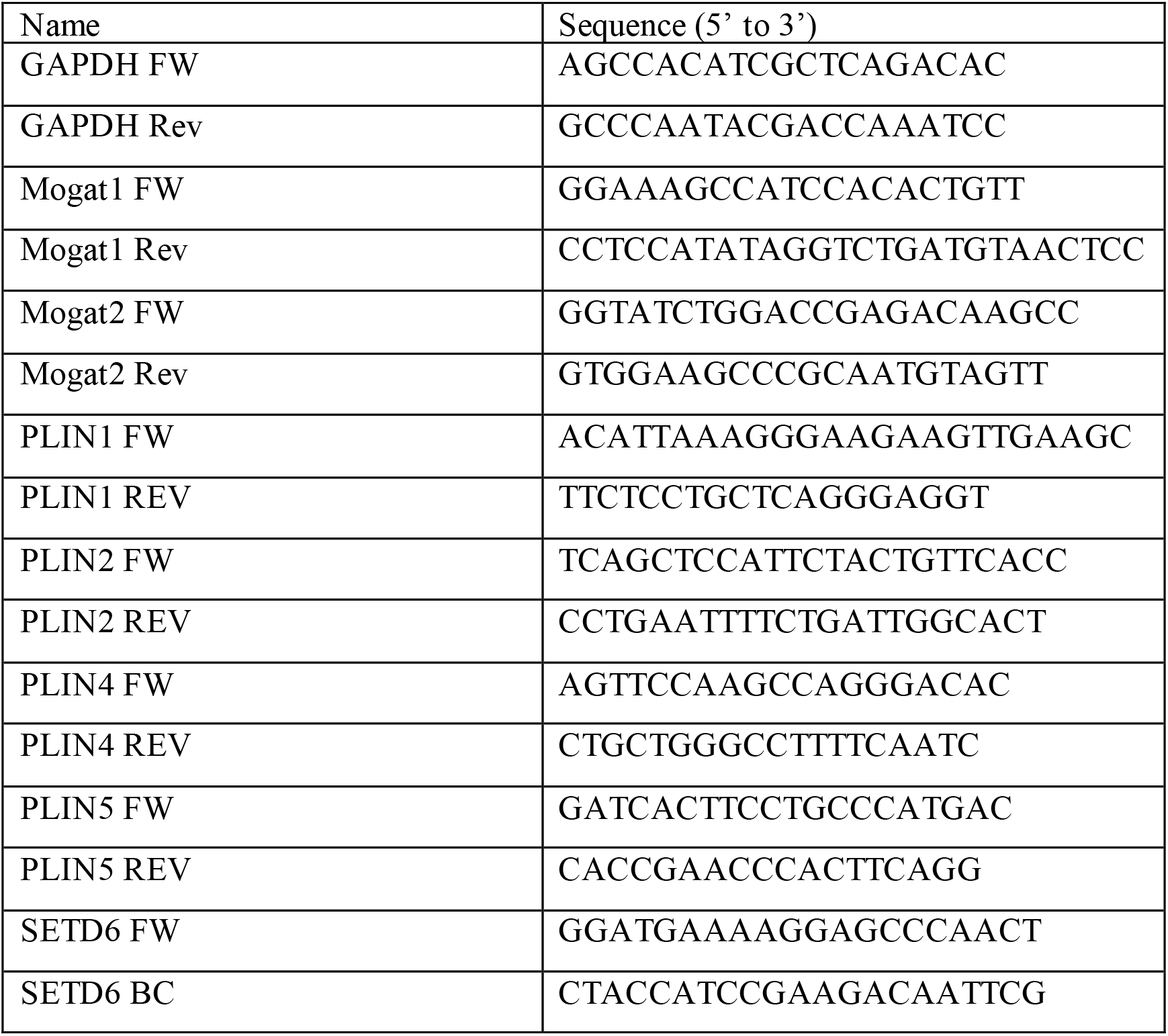
Primers for qPCR.

### Transfection

Cells were plated in 10-cm plates, and transfection was performed using polyethyleneimine (PEI) reagent (Polyscience Inc., 23966) for HEK293T cells or Mirus reagents; TransIT-X2 or TransIT-LT1 for HepG2 and Lipofectamine 2000 (Thermo fisher scientific) or TransIT-LT1 for HeLA cells, according to the manufacturer ’s instructions. For stable transfection in HepG2 cell lines, retroviruses were produced by transfecting HEK293T cells with the indicated pLenti constructs (empty, Flag PPARγ2 wild-type or Flag PPARγ2 K170R) and with plasmids encoding PMD2 and PX3. Target cells were infected with the viral supernatants and selected with 4 μg/ml puromycin.

### SETD6 knock-out by CRISPR/CAS9

HepG2 CRISPR/Cas9 SETD6 knock-out cells were generated using three independent sgRNAs targeting SETD6 cloned into the lentiCRISPR plasmid (Addgene, #49535). Cells transfected with the empty lentiCRISPR vector served as CRISPR control (CT) cells. Briefly, 3 × 10^5 HepG2 cells were plated per well in 6-well plates and transfected using TransIT-X2 reagent (Mirus) according to the manufacturer’s instructions. Following transduction and puromycin selection, single clones were isolated and expanded. Clones were validated by sequencing.

### Recombinant protein expression and purification

Escherichia coli BL21 transformed with a plasmid expressing a protein of interest were grown in LB media. Bacteria were harvested by centrifugation after IPTG induction and homogenized with ice-cold lysis buffer containing phosphate-buffered saline (PBS), 10mM imidazole, 0.1% Triton X-100, 1mM PMSF, and one complete mini protease inhibitors tablet (Roche). After adding 0.25mg/ml lysozyme for 30 min, the lysates were subjected to sonication on ice (18% amplitude, 1 min total, 10 sec ON/OFF). The tagged fusion proteins were purified on a His-Trap column using AKTA Pure protein purification system (GE). Eluted with 0.5M imidazole in PBS buffer, followed by overnight dialysis (PBS, 10% glycerol). Recombinant GST-SETD6 was overexpressed and purified from insect cells as previously described in (18) .

### In vitro methylation assay

The methylation assay reaction (total volume of 25 μl) contained 2 μg of His-Sumo mPPARγ2 wt, or K170R mutant proteins and 2 μg His-SETD6, 2mCi 3H-labeled S-adenosyl-methionine (SAM) (AdoMet, Amersham) and 5 μl of PKMT buffer (20mM Tris-HCl pH 8, 10% glycerol, 20mM KCl, 5mM MgCl2). The reaction tube was incubated overnight at 30°C. The reaction was resolved by SDS-PAGE for Coomassie staining (Expedeon, InstantBlueTM) and exposed to autoradiogram. For the non-radioactive (cold) methylation assay, 300 μM non-radioactive SAM was added (Abcam, ab142221).

### Enzyme-linked immunosorbent assay (ELISA)

2 μg of recombinant proteins (BSA, MBP-RelA, and His-Sumo mPPARγ2 wt) diluted in PBS were added to a sticky surface (Greiner Microlon) 96-well plate and incubated for 1 hr at RT followed by 3% BSA blocking in PBST for 1hr incubation at RT. After 3 washes with PBST, the plate was covered with 0.5 μg GST-SETD6 or GST protein (negative control) diluted in 1% BSA in PBST for 1 hr at RT. Plates were then washed and incubated with primary antibody (anti-GST, 1:4000 dilution) followed by incubation with HRP-conjugated secondary antibody (goat anti-rabbit, 1:2000 dilution) for 1 hr. Finally, TMB reagent and then 1N H2SO4 were added; the absorbance at 450 nm was detected using a Tecan Infinite M200 plate reader. Results are represented as relative absorbance fold compared to GST or BSA\ His-Sumo.

### Mass spectrometry analysis

*In vitro* methylation reaction was performed as described above, but instead of using a radioactive SAM, 300 μM non-radioactive SAM was added (Abcam, ab142221). Reactions were then sent for mass spectrometry analysis at the Weizmann Institute of Science as described previously (41).

### Antibodies and Western blot Analysis

Samples were denatured with Laemmli sample buffer (250mM Tris-HCl pH 6.8, 10% SDS, 30% glycerol, 5% β- mercaptoethanol, a pinch of bromophenolblue), boiled for 5 min at 95°C, resolved on 8%-12% SDS-PAGE gel, and then blotted onto PVDF membranes. Primary antibodies used were anti-Flag (Sigma, F1804), anti-HA (Millipore, 05–904), anti-actin (Abcam, ab3280), anti-SETD6 (Genetex, GTX629891), anti-PPARγ (CST, 2435), anti-PPARγ2 (Abcam, ab45036), anti-histone H3 (Abcam, ab10799), anti-PPARγ K170me1 (Abmart Inc.), and anti-pan-methyl lysine (Abcam, ab23366; used at 1:500 dilution for western blot analysis and 2 μg per reaction for immunoprecipitation experiments). HRP-conjugated secondary antibodies, goat anti-rabbit, goat anti-mouse, mouse anti-Rabbit were purchased from Jackson ImmunoResearch (111-035-144, 115-035-062, 211-032-171 respectively). For endogenic immunoprecipitation of the SETD6 experiment, HRP Goat anti-mouse light chain antibody (Jackson, 115-035-174) was used. Antibodies were diluted and prepared in TBST with 5% BSA or in PBST with 10% skim milk, according to the manufacturer ’s recommendations. The immobilized HRP antibodies were detected by ECL (Biological Industries). For Western blot analysis, cells were homogenized and lysed in RIPA buffer (50 mM Tris-HCl pH 8, 150 mM NaCl, 1% Nonidet P-40, 0.5% sodium deoxycholate, 0.1% SDS, 1 mM DTT, and 1:100 protease inhibitor mixture (Sigma)). Samples were resolved on SDS-PAGE, followed by Western blot analysis.

### Chromatin Extraction

10cm plate of HepG2 cells washed three times with PBSx1, harvested, and centrifuged (1,800*xg*, 5 min, 4°C). Firstly, the pellet was resuspended with 500ul of Buffer A (10mM Hepes pH 7.9, 10mM KCl, 1.5mM MgCl2, 0.34M Sucrose, and 10% Glycerol) supplemented with 0.1% Triton, 1Mm DTT, 1:200 Protease Inhibitor cocktail (PI), 100Nm PMSF and incubated on ice for 8 minutes. After another centrifugation, the supernatant was removed, and the pellet that consists mainly of intact nuclei was rewashed with Buffer A with the same supplementation without the addition of Triton. Then, the washed pellet was resuspended with 500ul of No Salt Buffer (3mM EDTA, 0.2mM EGTA) supplemented with 1:200 PI cocktail (P8340, Sigma) and 1mM DTT and kept on ice for 30 minutes with occasional vortexing. After centrifugation and removal of soluble nuclear proteins from the pellet, the pellet will consist of a chromatin fraction. Two different approaches were carried out to extract the soluble chromatin proteins; in the first one, the chromatin pellet was resuspended with 200 ul of Buffer A supplemented with 1:200 PI cocktail (P8340, Sigma) and 1:200 Benzonase (E1014-25KU, Sigma), incubated for 15 minutes at 37°C to solubilize the pellet. The second approach included resuspension of the chromatin pellet with 100 ul of Buffer A with the addition of 1:200 PI and then sonicated (Bioruptor, Diagenode) at high power settings for 3 cycles, 6 min each (30 sec ON/OFF). The soluble chromatin fraction was collected by low-speed centrifugation (1,800*xg*, 5 min, 4°C).

### Immunoprecipitation (IP)

Preclear of the chromatin fraction for 1 hr at RT was conducted with Magnetic CHIP protein A/G magnetic beads (Millipore, 16-663). Afterward, the soluble chromatin fraction was incubated overnight at 4°C with pre-conjugated A/G magnetic beads conjugated to the antibody of interest, washed three times, and analyzed by Western blot.

### Dual-Luciferase Assay

HepG2 cells were seeded in 24-well plates and transiently transfected with 0.1 μg Flag-PPARγ WT or K117R mutant, 0.1 μg firefly luciferase plasmid, containing *SETD6* promoter, and 0.1 μg Renilla luciferase plasmid. The total amount of transfected DNA in each dish was kept constant by the addition of empty vector, as necessary. Cell extracts were prepared 24h after transfection, and firefly luciferase activity was measured with the Dual-Glo Luciferase Assay system (Promega) and normalized to that of Renilla luciferase.

### Proximity Ligation Assay (PLA) image processing and data analysis

Cells were cultivated on coated coverslips, washed with PBS and fixed in cold 4% Paraformaldehyde (PFA) at RT for 15min. Cell permeabilization was performed using 0.5% Triton X-100 in PBS for 10min in RT. PLA (Duolink) was performed according to the manufacturer’s instructions (Sigma) using antibodies against GFP (Abcam, ab290) and Flag (Sigma, F1804) overnight in 4°C. Images were acquired by confocal Spinning disk microscopy with a 63x objective. The PLA units were calculated per cell as the ratio between the number of dots within the nucleus and the nucleus area (stained with DAPI, using Duolink mounting media). Only nuclei GFP stained were calculated (indicating successful GFP-SETD6 transfection). Each nucleus is then represented as a point in the quantification graph. Each frame represents maximum intensity projection for 40-60 Z-stacks captured. Image processing and data analysis was previously described in (51).

### RNA-Sequencing

Total RNA from HepG2 cells was extracted using the NucleoSpin RNA Kit (Macherey - Nagel) according to the manufacturer’s instructions. Samples were prepared in two biological replicates for control cells, four biological replicates for SETD6 KO and four biological replicas for PPARγ and PPARγ K170R mutant. Libraries were prepared using the INCPM-mRNA-seq protocol. Briefly, the polyA fraction (mRNA) was purified from 500 ng of total input RNA followed by fragmentation and the generation of double-stranded cDNA. Afterward, an Agencourt Ampure XP beads cleanup (Beckman Coulter), end repair, Abase addition, adapter ligation, and PCR amplification steps were performed. Libraries were quantified by Qubit (Thermo Fisher Scientific) and TapeStation (Agilent). The sequencing was done on a HiSeq instrument (Illumina).

### RNA-Sequencing analysis

The analysis of the raw sequence reads was carried out using the NeatSeq-Flow platform (https://doi.org/10.1101/173005). The sequences were quality trimmed and filtered using Trim Galore (version 0.4.5) (quality cutoff=25, length cutoff=25) and cutadapt (version 1.15) (DOI: https://doi.org/10.14806/ej.17.1.200). Alignment of the reads to the human genome (GRCh38) was done with RSEM (version 1.2.28) (Li and Dewey, 2011) (option ‘bowtie2’) and calculation of the number of reads per gene per sample was also done with RSEM. Quality assessment of the process was carried out using FASTQC (version 0.11.8) and MultiQC (version 1.0.dev0) (Ewels et al, 2016). Genes with low expression values (mean count < 1 over all samples) were excluded from differential expression analysis. Read counts for differential gene expression were analyzed with the DESeq2 R package (52) using the the NeatSeq-Flow platform DESeq2 module. Genes with fold change ≥ 1 and adjusted P < .05 were considered as significantly differentially expressed genes. Significant genes were clustered using the ‘eclust’ function from the factorextra R package (10.32614/CRAN.package.factoextra) with the default restriction of maximum clusters. In some cases, batch effect was included in the design and also corrected for visualization purposes using the sva R package (53). SVA: Surrogate Variable Analysis. R package version 3.52.0). For treated mutant PPAR<u>γ</u> vs. treated WT PPAR<u>γ</u> and for treated WT PPAR<u>γ</u> vs. EMPTY PPAR<u>γ</u>, genes with fold change ≥ 1 and adjusted P < .05. Enrichment analysis for KEGG pathways was performed using Enrichr (54,55). RNA seq data were deposited into the Gene Expression Omnibus database under accession number GSE328466.

### Real-Time qPCR

200 ng of the extracted RNA was reverse transcribed to cDNA using the iScript cDNA Synthesis Kit (Bio-Rad) according to the manufacturer’s instructions. Real-time qPCR was performed using the Light cycler 480 SYBR green master (Roche, 04-887-352). All samples were amplified in triplicates in a 384-well plate using the following cycling conditions: 5 min at 95°C, 45 cycles of amplification; 10 sec at 95°C, 10 sec at 60°C, and 10 sec at 72°C, followed by melting curve acquisition; 5 sec at 95°C, 1 min at 65°C and monitoring up to 97°C, and finally cooling for 30 sec at 40°C Gene expression levels were normalized relative to GAPDH gene and controls of the experiment.

### ChIP-qPCR

For DNA ChIP qPCR, DNA ChIP protocol was used from (56) with the following modifications: HepG2 Cells from 10 cm plate were cross-linked using 1.5% formaldehyde (Sigma, F8775) on a shaking platform for 10 min at room temperature. The cross-linking reaction was stopped by adding 1.25M glycine pH 8.5 for 5 min at room temperature. Cells were harvested and washed three times with cold phosphate-buffered saline, once with 2.5 ml of buffer I (10 mM HEPES [pH 6.5], 10 mM EDTA, 0.5 mM EGTA, 0.25% Triton X-100), and once with buffer II (10 mM HEPES [pH 6.5], 1 mM EDTA, 0.5 mM EGTA, 200 mM NaCl). Afterward, cells were resuspended with 500ul of lysis buffer (50mM Tris pH 8, 10mM EDTA, 1% SDS, 1:100 Protease Inhibitor cocktail), then sonicated (Bioruptor, Diagenode) at high power settings for 6 cycles, 6 min each (30 sec ON/OFF). The chromatin extract was then clarified by centrifugation for 15 min, full speed at 4°C, and diluted 5x in dilution buffer (20 mM Tris pH 8, 2 mM EDTA, 150 mM NaCl, 1.84% Triton X-100, and 0.2% SDS). Chromatin immunoprecipitation was performed as previously described (56,57). Chromatin was precleared overnight at 4°C with Magnetic CHIP protein A/G magnetic beads (Millipore, 16-663). The precleared sample was then immunoprecipitated in dilution buffer with Anti FLAG magnetic beads (Sigma, M8823) or Magnetic CHIP protein A/G magnetic beads (Millipore, 16-663). The immunoprecipitated complexes were washed once with TSE150 buffer (20 mM Tris pH 8, 2 mM EDTA, 1% Triton X-100, 0.1% SDS and 150 mM NaCl), TSE500 buffer (20 mM Tris pH 8, 2 mM EDTA, 1% Triton X-100, 0.1% SDS and 500 mM NaCl) buffer 3 (250 mM LiCl, 10 mM Tris pH 8, 1 mM EDTA, 1% sodium deoxycholate and 1% Nonidet P-40), and twice with TE buffer (10 mM Tris pH 8 and 1 mM EDTA). DNA was eluted with elution buffer (50 mM NaHCO3, 140 mM NaCl and 1% SDS) containing 0.2 μg/μl RNase A and 0.2 μg/μl Proteinase K. Finally, the DNA eluates were decross-linked at 65°C overnight with shaking at 900 rpm and purified by NucleoSpin Gel and PCR Clean-up kit (Macherey-Nagel), according to manufacturer’s instructions. Purified DNA was subjected to qPCR using specific primers (Table 3). Primers were designed based on PPARγ occupancy as previously described on PPARγ target gene (58,59). qPCR was performed using SYBR Green I Master (Roche) in a LightCycler 480 System (Roche). All samples were amplified in five replicates in a 384-well plate using the following cycling conditions: 5 min at 95°C, 45 cycles of amplification; 10 sec at 95°C, 10 sec at 60°C, and 10 sec at 72°C, followed by melting curve acquisition; 5 sec at 95°C, 1 min at 65°C and monitoring up to 97°C, and finally cooling for 30 sec at 40°C. The results were normalized to input DNA and presented as % input.

**Table 3.**
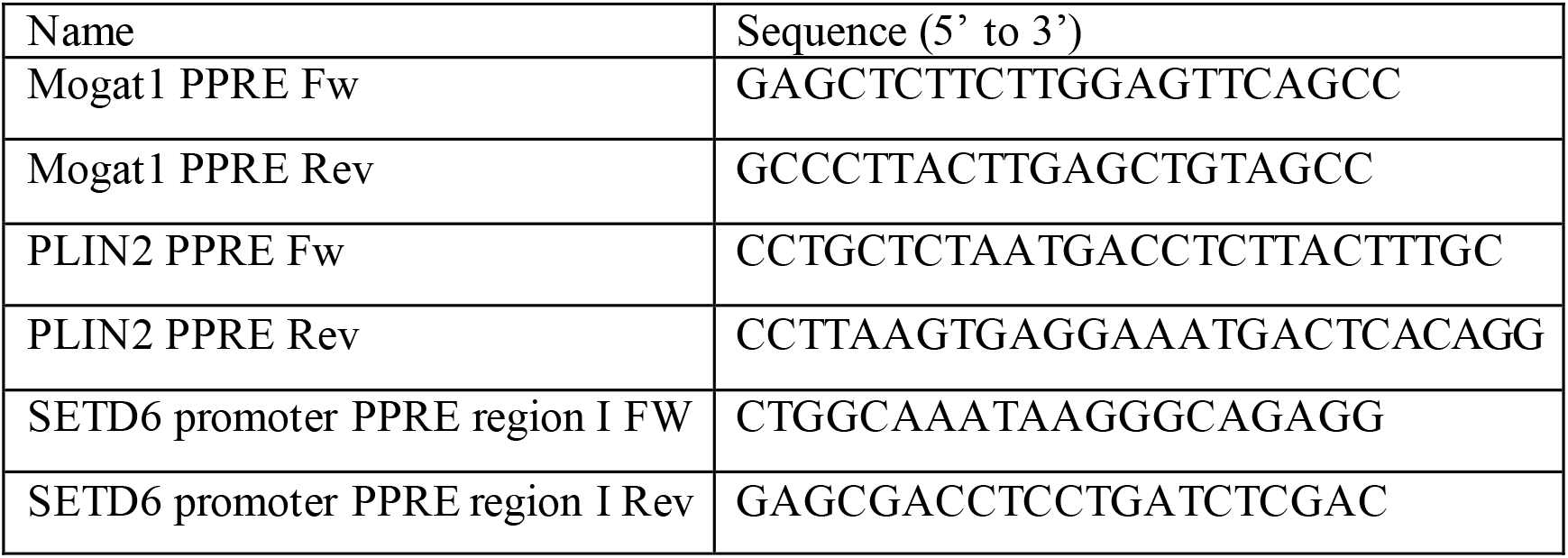
Primers for ChIP – qPCR.

### Structural modeling

Structural analyses were performed using the published crystal structure of the PPARγ–RXRα heterodimer bound to DNA and NCOA2 peptide (PDB: 3DZY). Structural visualization, residue modification, and analysis were carried out using UCSF ChimeraX (version 1.12). Mono-methylation of PPARγ K170 and phosphorylation of T166 were modeled using the “swapaa” command to generate the corresponding modified amino acid residues (M3L and TPO, respectively). Sidechain rotamers were selected using the ChimeraX rotamer library, and multiple rotamers were evaluated for K170me1. Potential steric clashes were assessed using the ChimeraX clashes/contacts tool with default parameters.

The structural model of the SETD6–PPARγ complex was generated using AlphaFold3. The predicted complex was visualized in ChimeraX, and the region surrounding the SET domain of SETD6 and the DNA-binding domain of PPARγ was analyzed. Structural models were used solely to generate mechanistic hypotheses and to visualize potential structural consequences of post-translational modifications; no quantitative energetic or molecular dynamics analyses were performed.

### Lipid Droplet Dynamics with Live Imaging Microscopy

20× 10^3^ cells of HepG2 cells were plated in 48 plate-well. On the next day, a live imaging microscopy was conducted for 20 or 24 hours to measure Oleic Acid (OA) accumulation. For Lipid Droplets biogenesis, cells were treated with 600 μmol/l of oleic acid [(OA, conjugated to albumin at 5:1 mol/l ratio), Sigma-Aldrich, St. Louis, MO, USA], with 200 ng/ml BODIPY 493/503 (D3922; Invitrogen, Carlsbad, CA, USA) for neutral lipid staining and 1 μg/ml Hoechst 33342 (H1399; Thermo Fisher Scientific, Inc., Waltham, MA, USA) was added for staining nuclei immediately before imaging. Imaging experiments were taken using a LionHeart FX (BioTek Instruments) configured with DAPI, GFP, and Brightfield light cubes. The fluorescence was measured with a 10x objective area scan with 3 beacons for each well and averaging the GFP signal for each well to account for variations in cell location in the well. Differences in cell number were corrected by using Hoechst fluorescence to normalize the BODIPY signal in each well. Data represent the mean of 3 wells per treatment. Image Analysis used primary and secondary masks of the captured digital images to determine the Mean GFP which represents the mean OA accumulation per cell. Primary mask analysis of the DAPI channel identifies individual cells by their nuclei. The secondary mask is denoted as the region up to 20 µm surrounding each nucleus and represents the cytoplasmic cell region. Scale bar indicates 200 µm.

### Statistical Analyses

Statistical analyses for all assays were analyzed with GraphPad Prism software, using Student’s two-tailed t-test (unpaired) when comparing two groups or one-way analysis of variance (ANOVA) with a Tukey’s *post hoc* test for comparing more than two groups. Two-way ANOVA was performed when two different categorical independent variables were tested on one dependent variable.

## Acknowledgments

This work was supported by grants to DL from The Israel Science Foundation (262/18 and 496/23), The Research Career Development Award from the Israel Cancer Research Fund and from the Israel Cancer Association. RZ is supported by The Israel Science Foundation (163/22).

## Author Contribution

N.N. and D.L. conceived and designed the study. N.N., D.G., and M.A. performed the majority of the experiments and analyzed the data. A.C. performed the mass spectrometry analysis. M.F. and TR performed the PLA and some of the IP experiments. Y.H., H.M., and A.R. contributed to data interpretation and provided conceptual input. TEB and RZ performed the structural modeling predictions. D.L. wrote the manuscript with input from all authors. All authors reviewed and approved the final version of the manuscript.

---

**Figure S1.**
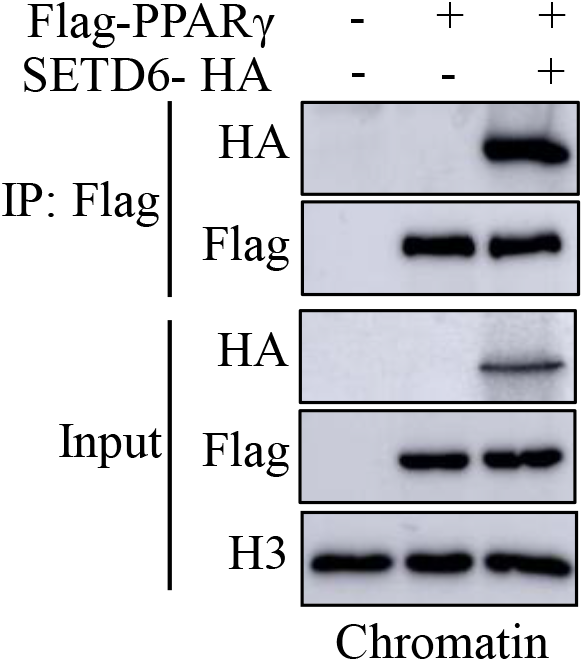
Interaction between overexpressed SETD6 and PPARγ in chromatin fractions. HEK293T cells were transfected with the indicated plasmids followed by immunoprecipitation with Flag antibody of the chromatin fraction. Samples were then subjected to WB analysis using the indicated antibodies.

**Figure S2:**
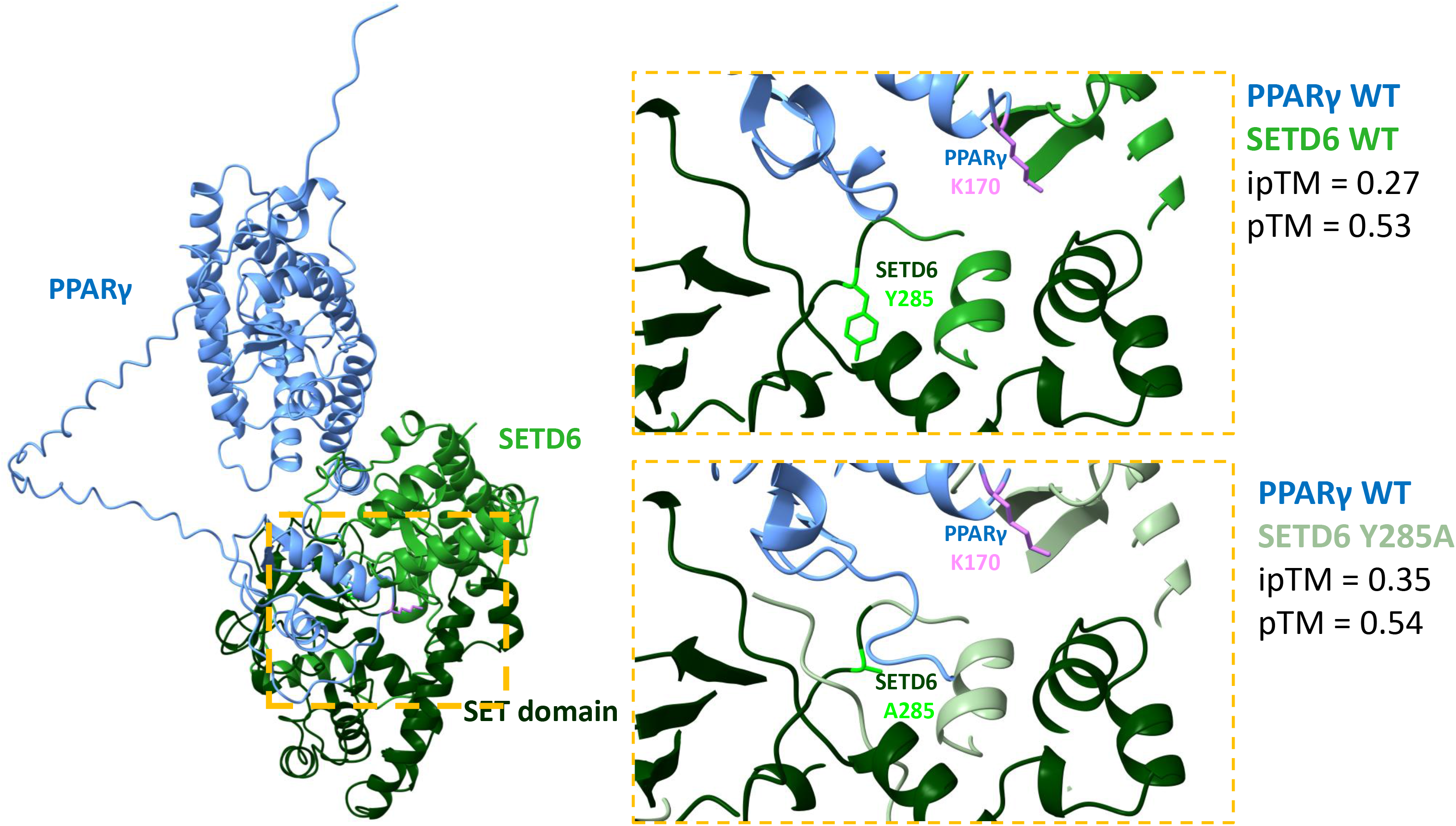
Structural modeling of the SETD6–PPARγ complex. **(A)** Structural model in cartoon representation of the predicted SETD6–PPARγ complex generated using AlphaFold. PPARγ is shown in blue, SETD6 is shown in green, and the SET domain is highlighted in dark green. Yellow boxes indicate the predicted interaction interface between the SET domain of SETD6 and the DNA-binding domain of PPARγ, shown at higher magnification in **(B)**. **(B)** Enlarged view of the predicted interaction interface comparing wild-type SETD6 (top) and the catalytically inactive mutant SETD6 Y285A (bottom). The catalytic residue Y285 (or A285 in the mutant) and PPARγ K170 are indicated in stick representation.

**Figure S3:**
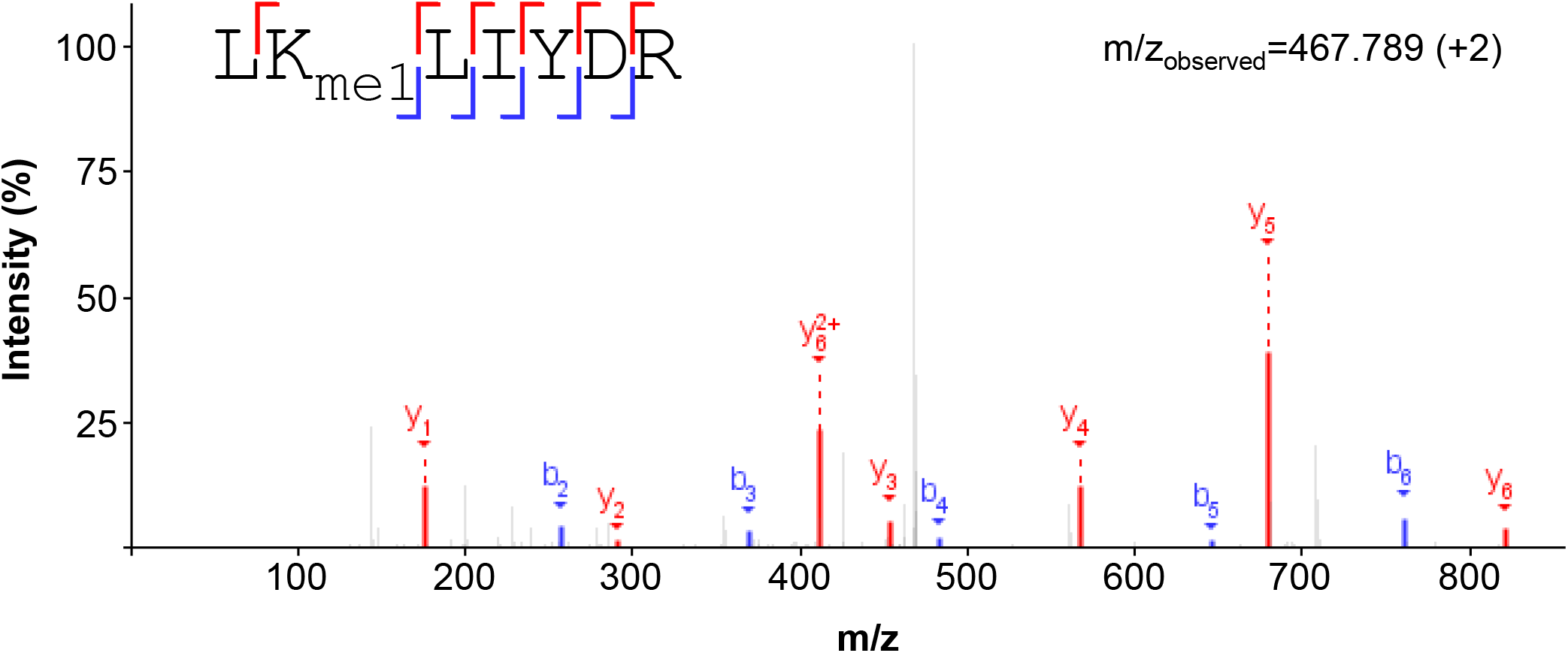
MS/MS spectra showing monomethylation of recombinant PPARγ at lysine-170 (LKme1LIYDR, m/zobserved =467.789 (z=+2)) after in vitro methylation by SETD6. The MS spectra was visualized with PDV software (v.2.0.0; (60)). The y-and b-ions are annotated and displayed as red, and blue, respectively.

**Fig S4:**
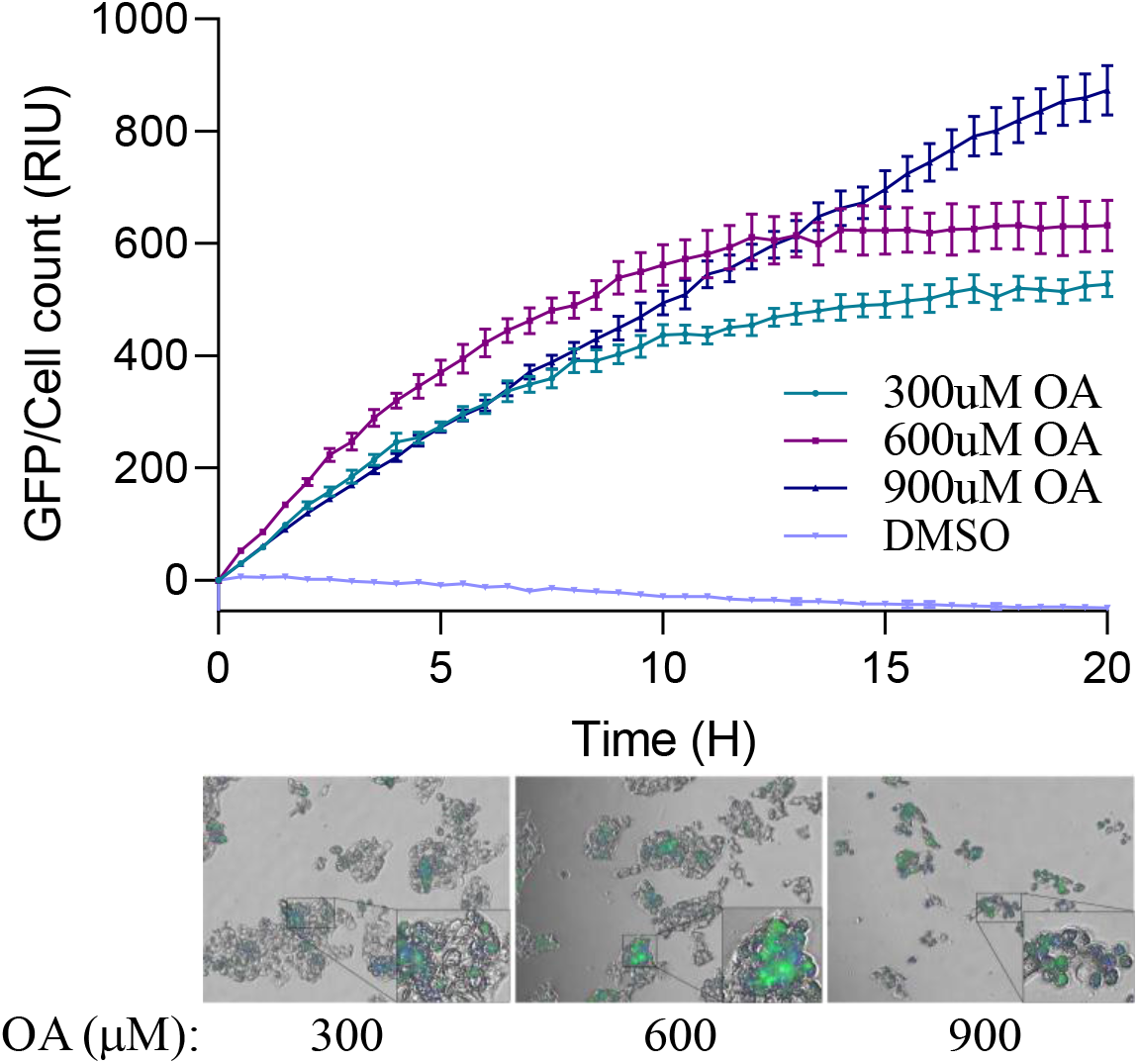
Top: OA accumulation curve of HepG2 cells over 20 hours challenged with 300 µM, 600 µM, and 900 µM of OA. 900 µM DMSO served as a control treatment. Mean green fluorescence was calculated as fluorescence signal divided by cell count. Data is analyzed from five beacons per well, with three wells per OA or DMSO treatment. Bottom- Representative images of last time point (20 H) with three concentrations of OA. For each image, a magnified area of interest is shown (black boxes).

**Fig S5:**
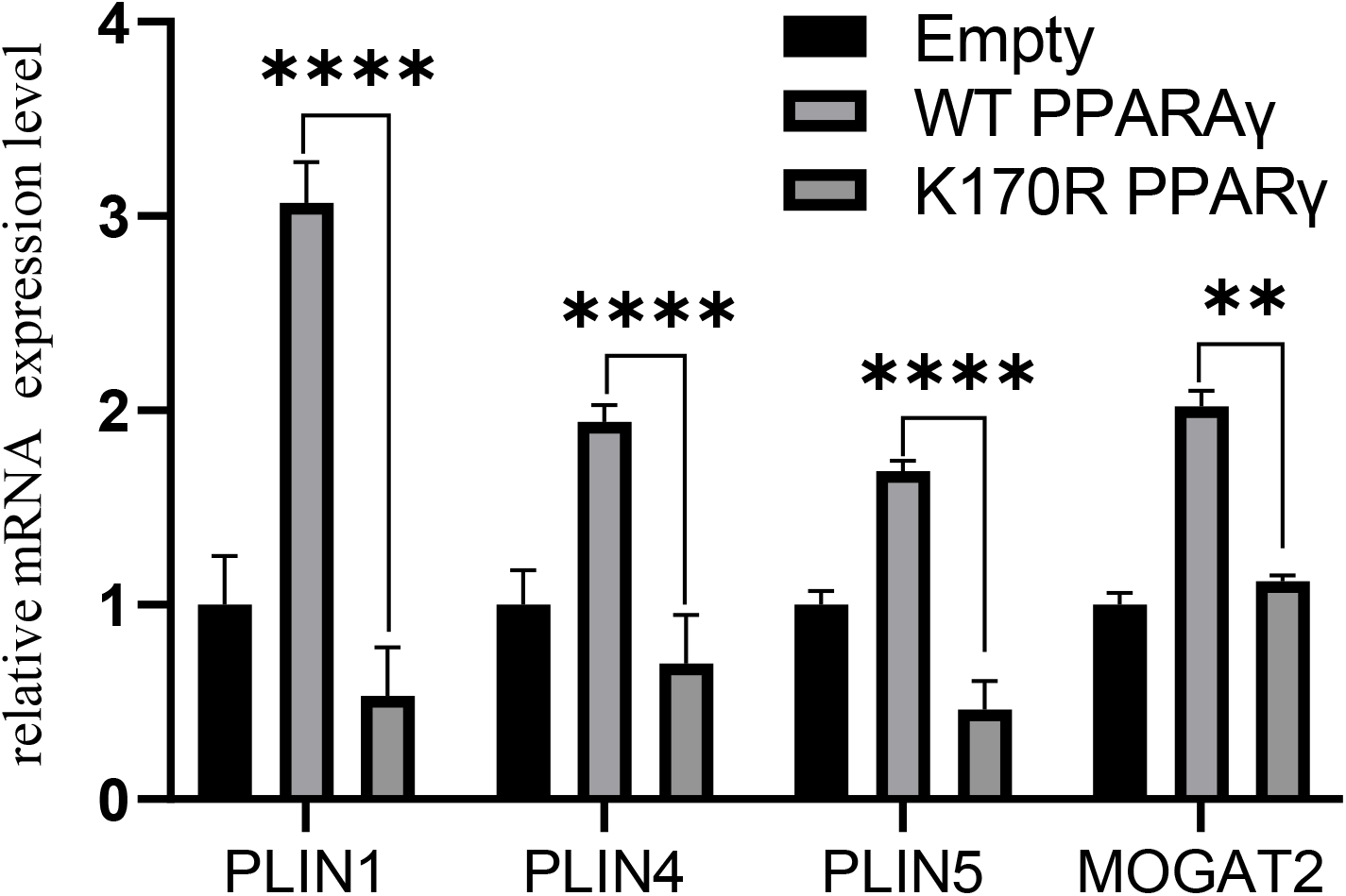
Transcript levels of the indicated genes were determined by qPCR of stably expressing HepG2 cells- Empty, Flag-PPARγ WT, and Flag-PPARγ K170R mutant. mRNA levels were normalized to GAPDH and then to Empty. error bars are S.E.M. *\*\*p* < 0.01; *\*\*\*\*p < 0.0001*.

**Figure S6.**
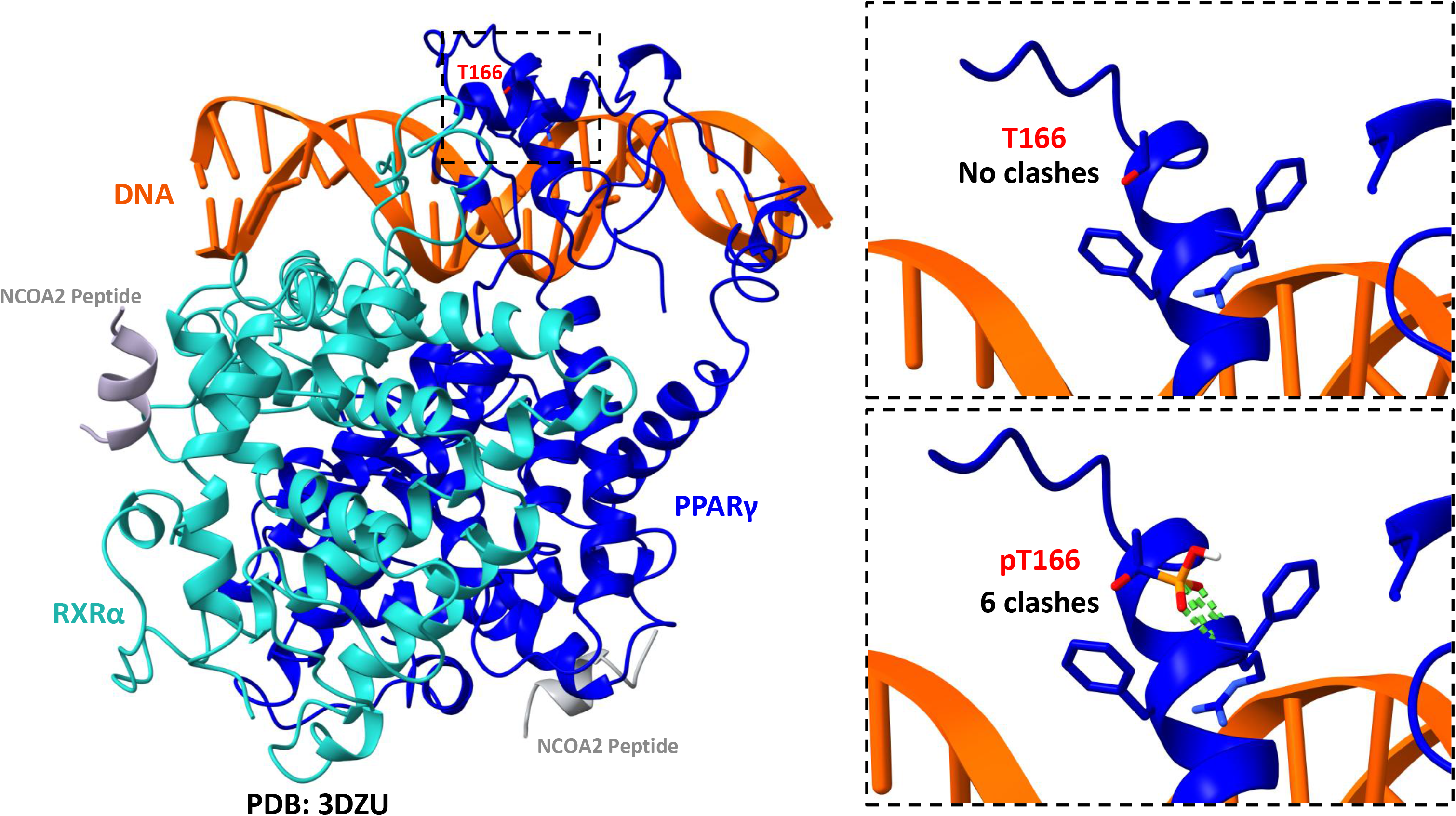
Structural modeling of T166 phosphorylation within the PPARγ DNA-binding domain. Structural model in cartoon representation of the PPARγ–RXRα heterodimer bound to DNA based on the published co-crystal structure (PDB: 3DZY). PPARγ is shown in blue, RXRα in cyan, DNA in orange, and NCOA2 peptides in gray. The boxed region highlights threonine 166 (T166) within the PPARγ DNA-binding domain. Enlarged views of the unmodified T166 residue (top) and the modeled phosphorylated T166 (pT166; bottom) are shown in stick representation. Predicted intramolecular steric clashes introduced by the phosphate group are indicated by green dashed lines.

